# Selective manipulation of excitatory and inhibitory neurons in top-down and bottom-up visual pathways using ultrasound stimulation

**DOI:** 10.1101/2025.02.14.638265

**Authors:** Yehhyun Jo, Xiaojia Liang, Hong Hanh Nguyen, Yeonseo Choi, Ga-Eun Bae, Yakdol Cho, Jiwan Woo, Hyunjoo Jenny Lee

**Author notes:** These authors contributed equally (Yehhyun Jo, Xiaojia Liang).

## Abstract

**Introduction:** Techniques for precise manipulation of neurons in specific neural pathways are crucial for excitatory/inhibitory (E/I) balance and investigation of complex brain circuits. Low-intensity focused ultrasound stimulation (LIFUS) has emerged as a promising tool for noninvasive deep-brain targeting at high spatial resolution. However, there is a lack of studies that extensively investigate the modulation of top-down and bottom-up corticothalamic circuits via selective manipulation of excitatory and inhibitory neurons. Here, a comprehensive methodology using electrophysiological recording and c-Fos staining is employed to demonstrate pulse repetition frequency (PRF)-dependent E/I selectivity of ultrasound stimulation in the top-down and bottom-up corticothalamic pathways of the visual circuit in rodents.

**Materials and methods:** Ultrasound stimulation at various PRFs is applied to either the lateral posterior nucleus of the thalamus (LP) or the primary visual cortex (V1), and multi-channel single-unit activity is recorded from the V1 using a silicon probe.

**Results and conclusion:** Our results demonstrate that high frequency PRFs, particularly at 3 kHz and 1 kHz, are effective at activating the bidirectional corticothalamic visual pathway. In addition, brain region-specific PRFs modulate E/I cortical signals, corticothalamic projections, and synaptic neurotransmission, which is imperative for circuit-specific applications and behavioral studies.

## Introduction

Precise modulation of specific neuronal circuits in the brain is critical for delineating network mechanisms and treating neuro-pathologies. In particular, effective regulation of excitatory-inhibitory (E/I) balance of top-down and bottom-up brain circuits is crucial for maintaining homeostasis and preventing neurological disorders^1–4^. The visual system is one of the most comprehensively mapped neural circuits in mammals, with well-established pathways that play crucial roles in linking key behavioral functions such as motor coordination, fight or flight instinct, memory, and decision-making^5–8^. This complex system is highly organized, with bidirectional corticocortical and corticothalamic pathways interacting to form a cohesive perception of the environment. These top-down and bottom-up processes are largely governed by the complementary dynamics of E/I neuronal projections across the corticothalamic pathways^9^. Thus, the ability to precisely and effectively manipulate E/I balance in the visual system using novel neuromodulation techniques is highly desired.

Among the widely implemented neuromodulation techniques, low-intensity focused ultrasound stimulation (LIFUS) has emerged as a promising method for noninvasive, deep-brain targeting with high spatial resolution^10–22^. There have been numerous studies that employ LIFUS in various brain regions including the visual cortex in rodents, non-human primates (NHP), and humans^13, 23–25^. However, despite enormous therapeutic potential, ultrasound is hindered by non-specific targeting of neurotransmitter release and neuronal types in the brain. Recently, there have been multiple reports that demonstrate the critical role of specific ultrasound waveforms in eliciting ultrasound-induced responses^14, 26–28^. Focused ultrasound beams are either generated as continuous waves at a set frequency (*f_0_*) or delivered in burst cycles at a specific pulse repetition frequency (PRF)^17, 29^. PRF-dependent activation of cortical pathways has been demonstrated using electrophysiological, calcium fiber photometry, and functional magnetic resonance imaging (fMRI) techniques^26, 28, 30, 31^. Furthermore, clinical studies have also shown selective GABA release in the cortical areas of humans^28^. While these results demonstrate the potential of LIFUS for selective manipulation of neuron types in disparate cortical areas, there is a lack of studies that show selective modulation of bidirectional E/I projections across corticothalamic regions and specific neurotransmitters involved in synaptic transmission. Thus, extensive investigations for developing precise LIFUS protocols are required for neuron-specific activation of selective pathways in the brain.

In this work, we selectively modulated E/I neurons of the corticocortical and corticothalamic pathways in the rodent visual system and established a region-dependent and waveform-dependent set of ultrasound parameters for precise circuit activation. To investigate effective LIFUS protocols for selectively driving E/I neurons in the visual cortex, we selectively targeted two key regions of the corticothalamic visual circuit with various PRFs and monitored single neuron activity from the visual cortex. In addition, brain-wide immunohistochemistry (IHC) using c-Fos co-stained with excitatory and inhibitory neurotransmitters was conducted to determine the effects of LIFUS on synaptic transmission. Our results demonstrate that PRF and region specificity of LIFUS play crucial roles in modulating the E/I balance of top-down and bottom-up pathways in the visual system. Functional neuron-type stimulation using an optimized LIFUS protocol enable precise modulation of bidirectional pathways within the circuit, which is crucial for noninvasive investigation of neural circuits and widespread therapeutic applications.

## Methods

### Animal care and housing

All animals and procedures used in the *in vivo* experiments were approved by the Institutional Animal Care and Use Committee (IACUC) at the Korea Advanced Institute of Science and Technology (KA2024-083-v1). A total of 43 male C57BL/6J and C57BL/6N mice (8∼ 12 weeks old) were used in this study. The mice were housed under a 12-h day/night cycle (lights on: 07:00, lights off: 19:00) with food and water provided *ad libitum*.

### Ultrasound stimulation and electrophysiological recording *in vivo* setup

All *in vivo* experiments were conducted in a dimly lit room with black-out curtains and red-light-filtered lamps. Mice were anesthetized and head-fixed on a rodent dual-arm stereotaxic apparatus (RWD Life Science Co., Ltd., China) using an isoflurane vaporizer system (RWD Life Science Co., Ltd., China). Isoflurane induction was at 4% and maintenance was at 1.5% at an O_2_ flow rate of 1 L/min. Vaseline gel and a soft fabric covering were applied to the eyes to maintain hydration and block light throughout the experiment (Unilever, NJ, USA). Lidocaine (2%) was subcutaneously injected into the skin to numb the area before surgery. Next, the skin and fascia overlying the top of the skull were carefully excised using sterilized surgical instruments, and the periosteum was gently removed using a povidone-iodine cotton swab. For precise stereotaxic ultrasound targeting and electrophysiological recording, the dorsal-ventral (DV) positions of bregma and lambda were calibrated to zero. A digital coordinate readout attached to the stereotaxic was used to pinpoint the areas directly above the thalamus (AP:-1.5 mm, ML:-1.5 mm) and primary visual cortex (AP:-3.6 to-5 mm, ML:-2.5 mm). Using a micro-drill (HP Bur, Saeshin, Republic of Korea), a craniotomy was performed to remove the skull over each labeled region. For LP stimulation, a 1.5-mm-diameter opening was created above the thalamus for ultrasound stimulation and a smaller 1-mm-diameter was created over the V1 for inserting the neural probe. For V1 stimulation, the 1.5-mm-diameter transducer opening was drilled over the V1, anterior to the 1-mm-diameter neural probe opening. The dura mater covering the exposed V1 was carefully removed using a 36G needle for probe insertion. During the surgical procedure, a medical-grade saline solution was used to hydrate the exposed brain to maintain physiological integrity and prevent desiccation.

Utilizing the left arm of a dual-arm stereotaxic system, we inserted either a 16 or 32-channel silicon neural probe (A1×16-5mm-25-177-CM16, A1×32-Poly2-5mm-50s-177-CM32, NeuroNexus, United States) into the visual cortex at an approximately 40-degree angle from the sagittal plane (AP:-5 mm, ML:-2.5 mm, DV:-1.2 mm). The electrodes spanned from the fourth to the sixth layer of the visual cortex. This was confirmed using a custom-written MATLAB program that utilized the Allen Brain Atlas with the insertion coordinates and angle to estimate the brain location^32^. The extracellular signals were acquired using an Intan RHD recording controller and Intan RHX data processing software (Intan Technologies, Los Angeles, CA, USA) with a bandwidth of 0.1 Hz to 15 kHz, sampling rate of 30 kHz, and 60 Hz notch filter. The recorded data was stored on a local computer drive for offline analysis.

Ultrasound stimulation was conducted using a focused ultrasound transducer (5 MHz, diameter = 24 mm, radius of curvature (ROC) = 38.1 mm; Hagisonic Inc., Korea) equipped with a 3D-printed collimator filled with degassed water and sealed with silicon elastomer (Kwik-Cast, World Precision Instruments, USA). The transducer was placed vertically over either the thalamic area or the visual cortical area using the right arm of the dual-arm stereotaxic system. Precise positioning of the ultrasound transducer was achieved by securing a needle to the 3D-printed collimator using a screw and calibrating the distance between the needle and the collimator tip using a caliper. The needle tip was used to pinpoint the precise location of the target location using bregma coordinates for LP (AP:-1.5 mm, ML:-1.5 mm, DV:-2.5 mm) and V1 (AP:-3.6 mm, ML:-2.5 mm, DV:-0.7 mm). Ultrasound gel was applied to fill the space between the collimator tip and the brain surface, serving as a coupling layer. We initiated ultrasound stimulation after a stabilization period lasting between one to one and a half hours, ensuring stable electrophysiological signals.

### Acoustic simulations of skull effects

Acoustic simulations for designing a skull-compensated acoustic lens were performed using a custom-written MATLAB code as reported in a previous paper^20^. Briefly, we incorporated 3D models of skulls from micro-computerized tomography (CT) imaging of *ex vivo* mouse skulls into the grid points of the 3D model mesh (Skyscan 1276, Bruker, Center for University-Wide Research Facilities, Jeonbuk National University). For simulations involving a cranial window, we removed the skull over the target region at the same position as the *in vivo* experiment setup (LP: AP:-1.5 mm, ML:-1.5 mm; V1: AP:-3.6 mm, ML:-2.5 mm). In the simulation setting, a transducer with a 12 mm radius was positioned at the same AP and ML coordinates as the target, maintaining a DV distance of 30 mm above the skull. The acoustic properties parameters were set to default values. To compare the effect of the skull on the ultrasound beam, we conducted 2D acoustic simulations with both an intact skull and a skull with a craniotomy. We applied the time-reversal recording method without considering the skull to match the beam on the target and reconstructed the beam. To evaluate the impact of the acoustic lens, we applied the time-reversal recording method while considering the skull and created an acoustic lens by compensating the recorded phase. We then reconstructed the beam with the lens on the transducer.

### Ultrasound beam profile measurement

To characterize the beam profile of our ultrasound transducer, we used a customized PVC water tank filled with degassed water, featuring a programmable XYZ motorized stage (Sciencetown Inc., Incheon, Republic of Korea) and a needle hydrophone (NH0500, Precision Acoustics, UK). To evaluate the effect of the craniotomy, a mouse skull with a cranial window (AP:-1.5 mm, ML:-1.5 mm) was placed between the collimator and hydrophone. A custom-made MATLAB program controlled the maneuvering of the hydrophone, enabling the scanning of a cubic volume measuring 10 mm × 1.5 mm × 1.5 mm centered at the focal spot of the ultrasound beam. The output of the hydrophone was amplified and DC-coupled (Precision Acoustics, UK), and the resultant signal was acquired by a digital oscilloscope (DSOX2022A, Agilent Technologies Inc.). The measured voltage data underwent conversion to peak acoustic pressure and was subsequently used to calculate spatial-peak pulse-average intensity (I_SPPA_) and spatial-peak temporal-average intensity (I_SPTA_). According to the sensitivity of the hydrophone (440 mV/MPa at 5 MHz), we obtained the peak acoustic pressure *P_0_* from the voltage readout. The acoustic intensity was then calculated using the formula: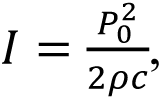 where ρ (density of water) = 1000 kg/m³ and *c* (speed of sound in water) = 1500 m/s. The lateral and axial resolutions of the ultrasound beam were determined by the full width at half maximum (FWHM) values.

### Ultrasound stimulation equipment and parameters

The ultrasound waveform was designed using two function generators (33220A, Agilent Technologies, CA, USA). The first function generator delivered a square wave at PRFs of 3 kHz, 1 kHz, and 80 Hz, corresponding to 3000 cycles, 1000 cycles, and 80 cycles, all at a 50% duty cycle. This PRF envelope waveform was triggered at 1-s bursts. The output of the first function generator was connected to the second function generator, which produced a sine wave tuned to the transducer’s resonance frequency of 5 MHz. The first function generator was controlled using a custom MATLAB program to generate 20-minute stimulation sessions with an inter-stimulation interval (ISI) of 10 seconds. The signal was amplified by an RF amplifier (240L, Electronics and Innovation, LTD, Rochester, USA) with a gain of 50 dB. The resultant peak pressure and I_SPPA_ were determined to be 51.25 W/cm^2^ and 1.24 MPa, respectively.

For the PRF parameters, three different PRF conditions of 3 kHz, 1 kHz, and 80 Hz were applied in a random sequence, each lasting 20 minutes. To account for potential confounds due to direct and indirect pathways between the auditory and visual cortices, a sham condition was implemented in which the ultrasound transducer was positioned away from the target location to create a high-impedance air gap between the collimator tip and the brain. For the control group, after craniotomy, neither ultrasound stimulation nor recording was performed.

### *In vivo* electrophysiology data analysis

To extract single-unit activity, spike sorting was performed using the open-source MATLAB-based automated spike-sorting software Kilosort 2.5^33^. We used the default parameters aside from modifying the threshold to [4,2]. The resulting cluster data were manually reviewed and refined using the open-source Python data analysis package Phy. The position of the electrodes within the visual cortex was verified using the Allen Brain Atlas and checked in real-time by observing the firing rate distribution across the cortical layers, as reported in previous studies^34, 35^. From the spike data, we manually selected isolated units based on the following criteria: less than 2% of spikes in the 2-ms long refractory period, normal spike waveform shapes, Gaussian distribution of amplitude, and peak-to-peak amplitude over 50 µV. Key features of the spike waveforms were used to distinguish between excitatory and inhibitory neurons. We separated single units into excitatory and inhibitory groups based on the trough-to-second peak latency and first peak-to-second peak ratio, calculated using a custom MATLAB code. Units with a trough-to-second peak latency greater than 0.7 ms were classified as fast-spiking units (FSUs, inhibitory neuron); otherwise, units were classified as regular-spiking units (RSUs, excitatory neuron). Among RSUs, units were further classified as RSU type 1 (RSU1) for first peak-to-second peak ratios lower than 1.5; otherwise, units were classified as RSU type 2 (RSU2).

Further analysis was conducted using the open-source MATLAB code available at https://github.com/cortex-lab/spikes (R2021b, MathWorks). Peri-stimulus time histograms (PSTH) were calculated with 10-ms bins spanning a 10-s interval, which included 5 s of both pre-and post-stimulation time to encompass baseline activity and potential long-term responses. Gaussian smoothing was applied with a standard deviation of 15 ms. To quantitatively compare the differences between different stimulation conditions, we analyzed the normalized firing rate, peak time, and decay time. The normalized firing rate was obtained by dividing the area under the PSTH curve during the 1-s stimulation by the area under the curve of the 1-s period prior to the stimulation. The peak time was defined as the time at which the firing rate reaches the maximum value within the 1-s stimulation period. The decay time was set as the time required for the firing rate to return to the baseline level before the stimulus was applied.

Analysis of local field potential (LFP) signals was conducted using the open-source MATLAB-based toolbox Brainstorm. The original data was downsampled to 1000 Hz and filtered using a bandpass filter from 0.5 Hz to 150 Hz. We defined the relevant epochs as segments ranging from-1000 ms to 2000 ms relative to the onset of each ultrasound stimulus. We conducted a time-frequency analysis (fast-Fourier transform) to calculate the average power for three physiologically critical frequency bands: delta (0.1 Hz to 4 Hz), theta (5 Hz to 7 Hz), and alpha (8 Hz to 12 Hz). The normalized power was obtained by dividing the area under the curve during the 1-s stimulation by the area under the curve of the 1-s period prior to the stimulation.

### Quantitative evaluation of ultrasound-induced thermal effects

To measure the temperature change in the brain due to ultrasound, we followed the same protocol as a previously reported paper^22^. Briefly, we employed a digital process controller (DP32PT, Omega Engineering, USA) connected to a duplex insulated T-type thermocouple wire (TT-T-36-SLE-50, Omega Engineering, USA) and inserted the probe tip into a brain-skull phantom. The system had an error margin of ±0.5°C and a tip size of 0.5 × 0.8 mm². For the brain phantom, a perfused and PFA-fixed mouse head with a large craniotomy centered at bregma was embedded in a 0.8% agarose gel. The thermocouple probe was inserted from the ventral-dorsal direction until the tip reached the edge of the brain (visible through the craniotomy). We aligned the focus of the ultrasound beam with the embedded thermocouple tip and then applied the same ultrasound protocol used in the *in vivo* LIFUS experiments. The temperature was measured at 5-s intervals for a total duration of 20 minutes and the measurement was repeated twice and the temperature values were averaged.

The mechanical index (MI) and thermal dose (CEM) were calculated using the following formulas:

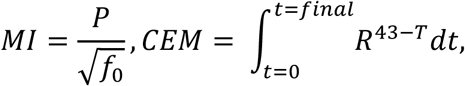

where *P* is the peak pressure, *f_0_* is the fundamental frequency, *t* is the time for the temperature change, and *T* is the temperature change. *T* was set at 38.5°C for a temperature change of 1.5°C and *R* was set at 0.25 for *T* < 43°C^36^.

### Histology and imaging

Post-stimulation, mice remained under isoflurane anesthesia (maintenance at 1∼2%) for 1 h 30 min before additional sedation with Avertin (20 μl/g, IP) and immediate transcardial perfusion using phosphate-buffered saline (PBS) followed by 4% paraformaldehyde (Sigma-Aldrich). The brains were carefully harvested and post-fixed in 4% paraformaldehyde at 4°C overnight. The samples were then washed in PBS and cryoprotected using 30% sucrose in PBS. Next, the brains were embedded in OCT (optimal cutting temperature) compound (Sakura) at-80°C before coronal sectioning in 50-µm-thick slices using a cryostat microtome (CM3050S, Leica). Free-floating sections were divided into two groups; the first group was washed in PBS before immediate incubation, while the second group was preserved at-20°C for long-term storage in a glycerol:ethylene glycol:PBS solution at a mixing ratio of 30:30:40.

For c-Fos antibody staining, brain sections were incubated with blocking and permeabilization solution (3% donkey serum and 0.3% Triton X-100 in PBS) for 1 h at room temperature. Primary antibody incubation was conducted with rabbit anti-Fos (ab190289, Abcam) diluted 1:500 in blocking solution at room temperature overnight. After a PBS washing session (3 × 15 min), sections were incubated in a secondary antibody of donkey anti-rabbit coupled to Alexa Fluor 488 (ab150073, Abcam) diluted 1:500 for 1 h at room temperature. Next, after another washing session in PBS (3 × 15 min), nuclear staining was conducted using DAPI diluted 1:20,000 in PBS, and the sections were washed again in PBS (2 × 10 min) to remove any excess staining solutions. Finally, the sections were mounted on glass slides (Thermo Scientific) with a mounting medium (Fluoromount), and a glass coverslip was applied for fluorescent imaging.

For c-Fos and vGLUT1 co-staining, brain sections were incubated with blocking and permeabilization solution (3% donkey serum and 0.3% Triton X-100 in PBS) for 1 h at room temperature. Primary antibody incubation was conducted with mouse anti-Fos (ab208942, Abcam) and rabbit anti-vGLUT1 (ab227805, Abcam) diluted 1:100 in blocking solution at room temperature overnight. After a PBS washing session (3 × 15 min), sections were incubated in secondary antibodies of donkey anti-mouse coupled to Alexa Fluor 594 (ab150108, Abcam) and donkey anti-rabbit coupled to Alexa Fluor 488 (ab150073, Abcam) diluted 1:500 for 1 h at room temperature. Subsequent steps were the same as the previous Fos-only staining.

For c-Fos and GABA co-staining, brain sections were incubated with blocking and permeabilization solution (3% donkey serum and 0.3% Triton X-100 in PBS) for 1 h at room temperature. Primary antibody incubation was conducted with rabbit anti-Fos (ab190289, Abcam) diluted 1:500 and mouse anti-GABA (ab86186, Abcam) diluted 1:100 in blocking solution at room temperature overnight. After a PBS washing session (3 × 15 min), sections were incubated in secondary antibodies of donkey anti-mouse coupled to Alexa Fluor 594 (ab150108, Abcam) diluted 1:200 and donkey anti-rabbit coupled to Alexa Fluor 488 (ab150073, Abcam) diluted 1:500 for 1 h at room temperature. Subsequent steps were the same as the previous Fos-only staining.

For hematoxylin and eosin (H&E) staining, immediately following cryosectioning, free-floating brain sections were mounted on Superfrost microscope slides (Sigma Aldrich) and left to dry at room temperature overnight. Then, the following protocol was used to stain the sections before applying mounting solution and a glass cover slip: 70% EtOH 2 min → DW 2 min → Hematoxylin 10 min → DW 2 min → 1% HCl 5 s → DW 2 min → 0.4% Ammonia 2 min → DW 2min → Eosin 2 min → 70% EtOH 2 min → 80% EtOH 2 min → 90% EtOH 2 min → 95% EtOH 2min → 100% EtOH 2 min → 100% EtOH 2 min → 100% EtOH 2 min → Xylene 2 min → Xylene 2 min → Xylene 2 min → Xylene 2 min.

All fluorescent sections were imaged using a confocal microscope (LSM780, Zeiss) or slide scanner (Pannoramic SCAN II, 3DHISTECH Ltd.) in the EM & Histology Core Facility, at the BioMedical Research Center, KAIST. Image analysis was conducted using QUINT workflow, following previously published methods^16, 21, 37–41^. Briefly, stain-positive cells were identified using Ilastik and Image J, and then brain sections were aligned to the Allen Mouse Brain Atlas using DeepSlice, QuickNII, and VisuAlign. For cell counting, Nutil Quantifier was used for semi-automated parsing of object data before analysis with a custom-written MATLAB program. A 3D meshview was used to visualize the stain-positive cells across various brain regions. Lastly, Image J was used to determine the ratio of Fos-positive cells to GABAergic and glutamatergic cells. For imaging of H&E-stained sections, a bright-field microscope was used (CX41, Olympus).

## Statistical analysis

Normal distribution of all *in vivo* data sets was confirmed using Shapiro-Wilk’s normality test. For electrophysiological data, one-way analysis of variance (ANOVA) with Tukey’s *post-hoc* test was conducted, using ultrasound stimulation conditions as a single factor. For immunohistology data, parametric analysis was conducted using a one or two-sided unpaired Student’s *t*-test, and one-way ANOVA with Tukey’s *post-hoc* test was conducted using ultrasound stimulation conditions as a single factor.

All data were analyzed using Microsoft Excel (Microsoft Co., WA, USA), OriginPro 2019 (OriginLab Co., MA, USA), and MATLAB R2022b (Mathworks, MA, USA). Neural spike sorting and neuronal type classification were conducted using the open-source program Kilosort 2.5. Unless stated otherwise, all data are presented as means ± standard error of means (s.e.m.) and a *p*-value of <0.05 was considered significant. For analysis of electrophysiological data, outliers were defined as data points exceeding the third quartile (3Q) by three times the interquartile range (IQR). Graphical illustrations of biological components in all relevant figures were created with Adobe Illustrator 2023 and BioRender.com.

## Results

### Overview of target brain circuit and stimulation protocol

There are two major corticothalamic pathways in the rodent visual system that are responsible for motor coordination, object recognition, and visual signal processing^42–47^. One pathway extends from the retina to the dorsal lateral geniculate nucleus (dLGN) and reaches the primary visual cortex (V1), while the other traverses from the retina to the superior colliculus (SC), which then travels to the lateral posterior nucleus (LP) and reaches the visual cortex, including higher-order processing regions such as the V2 (Fig. 1a). In this study, we developed a stimulation paradigm for the LP-V1 circuit by mechanically scanning the lateral thalamus with LIFUS and verifying acute responses using electrophysiological and IHC readouts. We found that direct dLGN stimulation using LIFUS in sedated mice did not generate significant responses, which is consistent with previous reports that showed inactivation of the dLGN-V1 circuit of rodents to ultrasound stimulation under anesthesia^48^. Using a 5-MHz focused transducer and 32-channel neural probe, we stimulated the (1) LP region while measuring single-unit neuronal activity from the V1 (LP-V1 group) and (2) V1 region while recording from the V1 (V1-V1 group) (Fig. 1b and Supplementary Fig. 1a-c). In both regions, we varied the PRFs, consisting of 3 kHz, 1 kHz, and 80 Hz waveforms (Fig. 1b and Supplementary Fig. 2a). This gamut of frequencies was determined based on physiologically critical high-frequency neuronal burst activities (3 kHz and 1 kHz) and low-frequency gamma oscillation (80 Hz), which is linked to higher order cognitive function^16, 17,27, 49–53^.

**Fig. 1.**
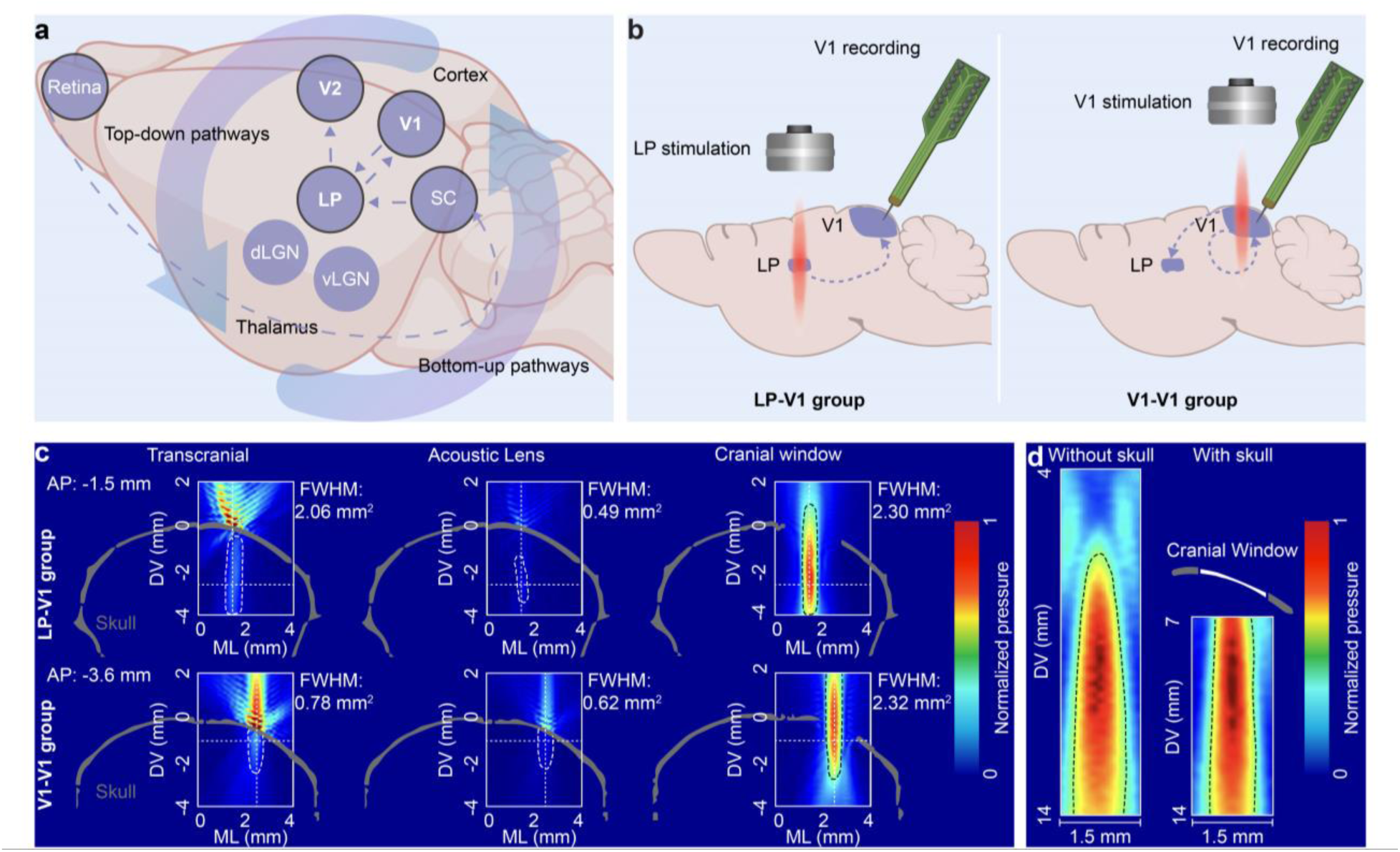
Experimental setup for LIFUS and electrophysiological recording in the visual pathway of rodents. **a**, Overview of the top-down and bottom-up pathways in the visual circuit of rodents. **b**, Schematic of the LIFUS and electrophysiological recording system. The ultrasound transducer was positioned vertically over either the LP (LP-V1 group) or V1 (V1-V1 group) and a silicon neural probe was inserted into the V1 at a 40-degree angle from the sagittal plane. The purple dashed line indicates the innervation pathways of the top-down and bottom-up corticothalamic and corticocortical circuits. **c**, Simulation of the ultrasound beam at the resonant frequency for the LP-V1 and V1-V1 groups with an intact skull (left), a skull-compensated acoustic lens (center), and a craniotomy (right). White dotted line indicates FWHM beam size. Color bar indicates normalized acoustic pressure. **d**, Beam profile measurement of the transducer at the resonant frequency without the skull (left) and with the craniotomized skull (right). Black dotted line indicates FWHM beam size. Color bar indicates normalized acoustic pressure.

For precise targeting of deep-brain regions using ultrasound, the skull effect must be considered. The beam aberration is particularly problematic for laterally located brain regions due to the large incident angle of the beam at the skull surface^16^. While there are skull-compensation methods using acoustic lenses and phased arrays, a craniotomy eliminates the skull issue, particularly for *in situ* acute experiments^20^. Using acoustic simulations, we demonstrated that a small craniotomy allowed the ultrasound beam to precisely target the LP and V1 without any attenuation and aberration (Fig. 1c, Methods). Our simulation consisted of a 5-MHz beam and a micro-CT skull model for LP and V1 targeting with: (1) an intact skull, (2) a skull-compensated acoustic lens, and (3) a craniotomy. For both the LP-V1 and V1-V1 groups, the full width at half maximum (FWHM) beam shape and the beam attenuation after skull transmission were significantly affected by the presence of the skull. Compared to the transcranial and acoustic lens cases, the craniotomy produced negligible beam aberration and attenuation. To experimentally verify our simulations, we measured the beam profile of the transducer without the skull and with a craniotomized skull (Fig. 1d and Supplementary Fig. 2b, c). The beam measured without the skull had a FWHM of 5.40 mm² and a focal point at DV = 9.00 mm from the collimator, while the beam with the craniotomized skull had a FWHM of 4.56 mm² in the brain and a focal point at DV = 1.95 mm from the collimator. These results demonstrate significant skull-induced beam attenuation and aberration at a fundamental frequency of 5 MHz, while the skull craniotomy provided a robust and reproducible beam without attenuation and aberration.

In this study, we maintained the acoustic intensity and stimulation duration the same across all experimental groups, with PRF as the only variable. At a stable level of anesthesia for the duration of the stimulation (isoflurane induction was at 4% and maintenance was at 1.5% at an O_2_ flow rate of 1 L/min), a spatial-peak, pulse-average intensity (I_SPPA_) of 51.25 W/cm^2^ (peak pressure of 1.24 MPa) was required to reliably activate the neural pathways (Supplementary Fig. 2d). While this is relatively high compared to other LIFUS reports, the mechanical index (MI) and thermal dose were 0.55 and 0.04 cumulative equivalent minutes (CEM), respectively, which are lower than the MI cavitation threshold of 1.9 and thermal dose threshold of 1 CEM^36^. To verify the biological safety for this level of intensity, we conducted a thermal test using a thermocouple and brain phantom, which demonstrated an instantaneous peak temperature change of approximately 1.27°C during a 1-s trigger over the 20-min total stimulation duration (Supplementary Fig. 3a-c). While this is higher than 1°C, considering the lack of a heat-dissipating perfusion mechanism in the phantom and the low MI and CEM values, we hypothesized that our protocol would not cause any tissue damage. We confirmed that there was no structural damage in the brain post-stimulation using bright-field images of hematoxylin and eosin (H&E)-stained brain sections (Supplementary Fig. 4a, b; Methods).

### LIFUS can bidirectionally activate the LP-V1 pathway

To verify the activation of a specific circuit within the complex visual system, we targeted either the LP or V1 at a PRF of 3 kHz and measured single neuron activity from the V1 for both cases. LIFUS was triggered for a burst duration of 1 s at an I_SPPA_ of 51.25 W/cm^2^ (peak pressure of 1.24 MPa). We first delivered LIFUS to the LP of anesthetized wild-type mice and recorded neural spikes from the V1 (LP-V1 group) (*n* = 11). Multi-unit activity (MUA) of neural spikes was observed in the raw signal recording and single-unit activity was extracted using an analysis pipeline based on Kilosort. We found a significant increase in the firing rate of neurons in the V1 during LIFUS of the LP compared to the sham condition (Fig. 2a, Supplementary Fig. 5, and Supplementary Fig. 6; Methods). LIFUS was delivered at time zero for a total stimulation duration of 1 s, with single-unit activity recorded across 9651 trials in the stim condition and 10865 trials in the sham condition. The elevated response was maintained through the 1-s stimulation, peaking at approximately 0.36 s after the trigger, indicating a mechanistic relationship between ultrasound neuromodulation duration and the refractory behavior of neurons (Fig. 2a).

**Fig. 2.**
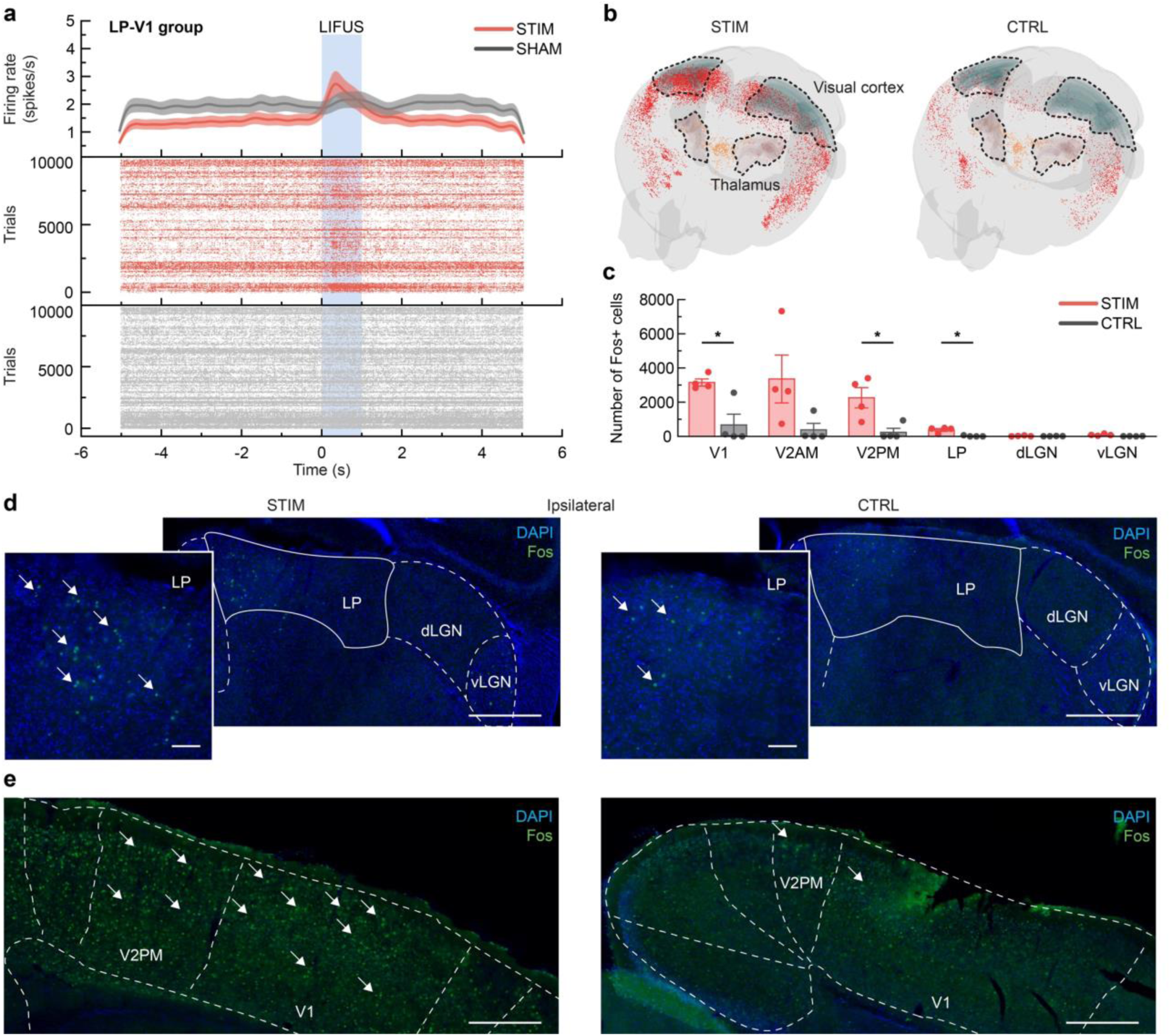
Neural activation of the LP-V1 group (Stimulation of LP and recording from V1). **a**, Peri-stimulus time histogram (PSTH, solid line: mean value, shaded area: standard error of the mean (SEM), bin size: 10 ms, Gaussian smoothing SD: 15 ms) and raster plots of single-unit spiking activity in the V1 for STIM and SHAM conditions. Blue shaded bar indicates the 1-s LIFUS duration. **b**, Representative 3D mesh views of Fos-positive cells in the thalamus and visual cortex for STIM and CTRL conditions. **c**, Number of Fos-positive cells in the V1, V2AM, V2PM, LP, dLGN, and vLGN for STIM and CTRL conditions. *p < 0.05, two-sided unpaired Student’s t-test, n = 8. **d**, Representative Fos IHC images of the ipsilateral thalamus for STIM and CTRL conditions. Scale bars, 100 µm (small), 400 µm (large). **e**, Representative Fos IHC images of the ipsilateral visual cortex for STIM and CTRL conditions. Scale bar, 400 µm.

We then performed LIFUS of the V1 while recording from the V1 to compare electrophysiological response dynamics between the corticothalamic (LP-V1 group) and corticocortical (V1-V1 group) activation pathways (*n* = 8). As expected, we observed a significant increase in the neuronal firing rate in the V1 during LIFUS compared to the sham condition (Fig. 3a). Stimulation was delivered for 1 s, with single-unit activity recorded across 5588 trials in the stim condition and 4221 trials in the sham condition. The response also remained elevated throughout the 1-s stimulation, peaking at approximately 0.35 s after the trigger delivery. The 10-ms difference in peak response time between the LP-V1 group and V1-V1 group confirmed the latency of neuronal signal transmission from the thalamus to the cortex^54^. These electrophysiological results demonstrate the capability of LIFUS to acutely modulate neural spiking activity of connected brain circuits.

**Fig. 3.**
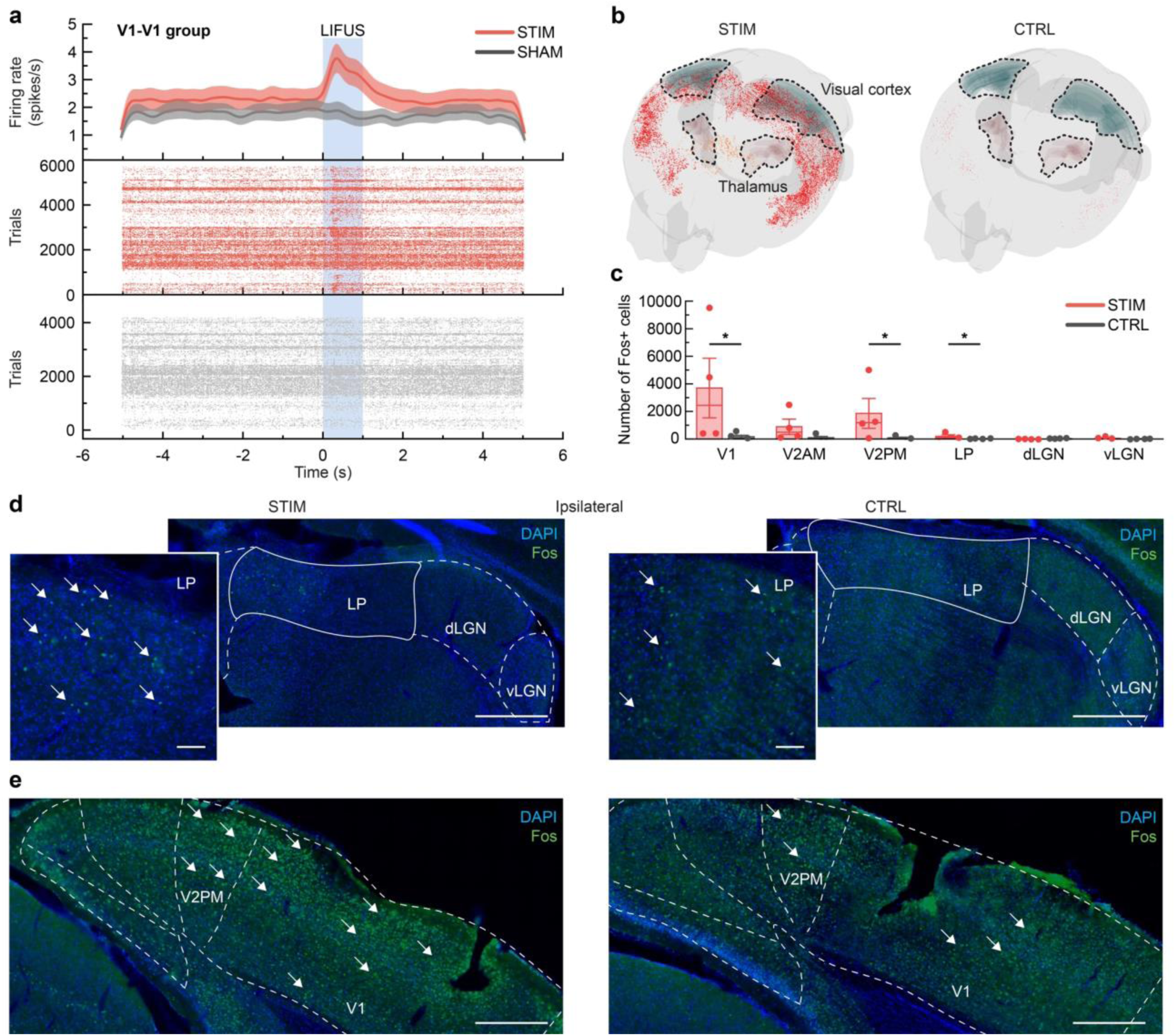
Neural activation of the V1-V1 group (stimulation of V1 and recording from V1). **a**, Peri-stimulus time histogram (PSTH, solid line: mean value, shaded area: standard error of the mean (SEM), bin size: 10 ms, Gaussian smoothing SD: 15 ms) and raster plots of single-unit spiking activity in the V1 for STIM and SHAM conditions. Blue shaded bar indicates the 1-s LIFUS duration. **b**, Representative 3D mesh views of Fos-positive cells in the thalamus and visual cortex for STIM and CTRL conditions. **c**, Number of Fos-positive cells in the V1, V2AM, V2PM, LP, dLGN, and vLGN for STIM and CTRL conditions. *p < 0.05, two-sided unpaired Student’s t-test, n = 9. **d**, Representative Fos IHC images of the ipsilateral thalamus for STIM and CTRL conditions. Scale bars, 100 µm (small), 400 µm (large). **e**, Representative Fos IHC images of the ipsilateral visual cortex for STIM and CTRL conditions. Scale bar, 400 µm.

To quantitatively analyze the network activation across brain regions, we conducted immunohistochemistry (IHC) of c-Fos, an immediate early gene (IEG) that is a well-established marker for identifying neuronal activity. The temporal activation of c-Fos provides a snapshot of neural dynamics in response to a specific internal or external stimulus such as LIFUS^55^. We measured the Fos-positive cell count in the LP, V1, anteromedial and posteromedial visual areas (V2AM and V2PM), dorsal and ventral lateral geniculate nucleus of the thalamus (dLGN and vLGN), central and basolateral amygdala (CeA and BmA), superior colliculus (SC), and auditory cortex (Aud) for the LP-V1 and V1-V1 groups. For each group, the LIFUS condition was compared to a control condition. Whole coronal brain sections were obtained and analyzed using a custom QUINT workflow to generate 3D visualizations of Fos-positive cells in the lateral thalamus and visual cortex (Supplementary Fig. 7; Methods). In the LP-V1 group, we found a significant increase of Fos-positive cells in the stimulated group compared to the control group for the LP, V1, V2PM, and Aud areas (Fig. 2b, c and Supplementary Fig. 8; two-tailed unpaired *t*-test, \**p* < 0.05, *n* = 8). In particular, the IHC results showed low levels of Fos-positive cells in the dLGN and vLGN regions, indicating that those visual pathways were not activated (Fig. 2c and Supplementary Fig. 9a, b; two-tailed unpaired *t*-test, *p* > 0.05, *n* = 8). Representative images of the contralateral and ipsilateral brain sections of the thalamus and visual cortex indicated that LIFUS activated both hemispheres of the brain, which is consistent with the bilateral connectivity of visual networks^56^ (Fig. 2d, e and Supplementary Fig. 9a, b). These images showed elevated levels of Fos-positive cells in the LP, V1, and V2 regions, while the dLGN and vLGN regions showed low levels of Fos activity. This verified precision activation of the LP-V1 neural circuit by selectively targeting the LP region using LIFUS.

Moreover, in the V1-V1 group, we also found a significant increase of Fos-positive cells in the stimulated group compared to the control group in the LP, V1, V2PM, BmA, and SC areas (Fig. 3b, c and Supplementary Fig. 10; two-tailed unpaired *t*-test, \**p* < 0.05, *n* = 9), along with low-level activation in the dLGN and vLGN (Fig. 3c and Supplementary Fig. 11a, b; two-tailed unpaired *t*-test, *p* > 0.05, *n* = 9). Both contralateral and ipsilateral areas of the thalamus and visual cortex were activated, indicating that LIFUS activated both hemispheres of the brain (Fig. 3d, e and Supplementary Fig. 11a, b). These representative images showed elevated levels of Fos-positive cells in the LP, V1, and V2 regions, while the dLGN and vLGN regions showed low levels of Fos activity. These IHC results were consistent for both the LP-V1 group and the V1-V1 group, indicating that LIFUS of either the LP or V1 at a PRF of 3 kHz selectively activated the LP-V1 circuit of the visual system.

To further analyze the bidirectional network effects of LIFUS on the LP-V1 circuit, we analyzed the neural firing rates in V1 cortical layers 4, 5, and 6 (L4, L5, and L6; Fig. 4a and Supplementary Fig. 12a, b, Methods). Representative single unit spikes from each of the four deep cortical layers demonstrated robust recording across L4, L5 and L6 (Fig. 4a and Supplementary Fig. 13). It is well established that the LP receives visual signals from layers 5 and 6 (L5 and L6) and strongly projects E/I signals to layers 1 and 5 (L1 and L5) of the V1^35, 45, 47, 56–59^. These innervation patterns suggest that E/I balance of the LP-V1 pathways is regulated by specific neuronal routes. As expected, we found an acute increase in the firing rate of neurons in V1 during LIFUS in L4, L5, and L6 for both the LP-V1 and V1-V1 groups (Fig. 4b-g). In the LP-V1 group, we observed the highest increase in firing rate in L5, which is consistent with well-established LP-to-V1 layer projections (Fig. 4c). In L4 and L6, the LIFUS-induced increase in firing rate was comparatively less pronounced (Fig. 4b, d). For the V1-V1 group, we found the greatest increase in firing rate in L4 and L5 of the V1, which indicates that LIFUS delivered at a PRF of 3 kHz effectively activated cortical neurons (Fig. 4e, f). In L6, the increase in firing rate was lower than that of L4 and L5 (Fig. 4g).

**Fig. 4.**
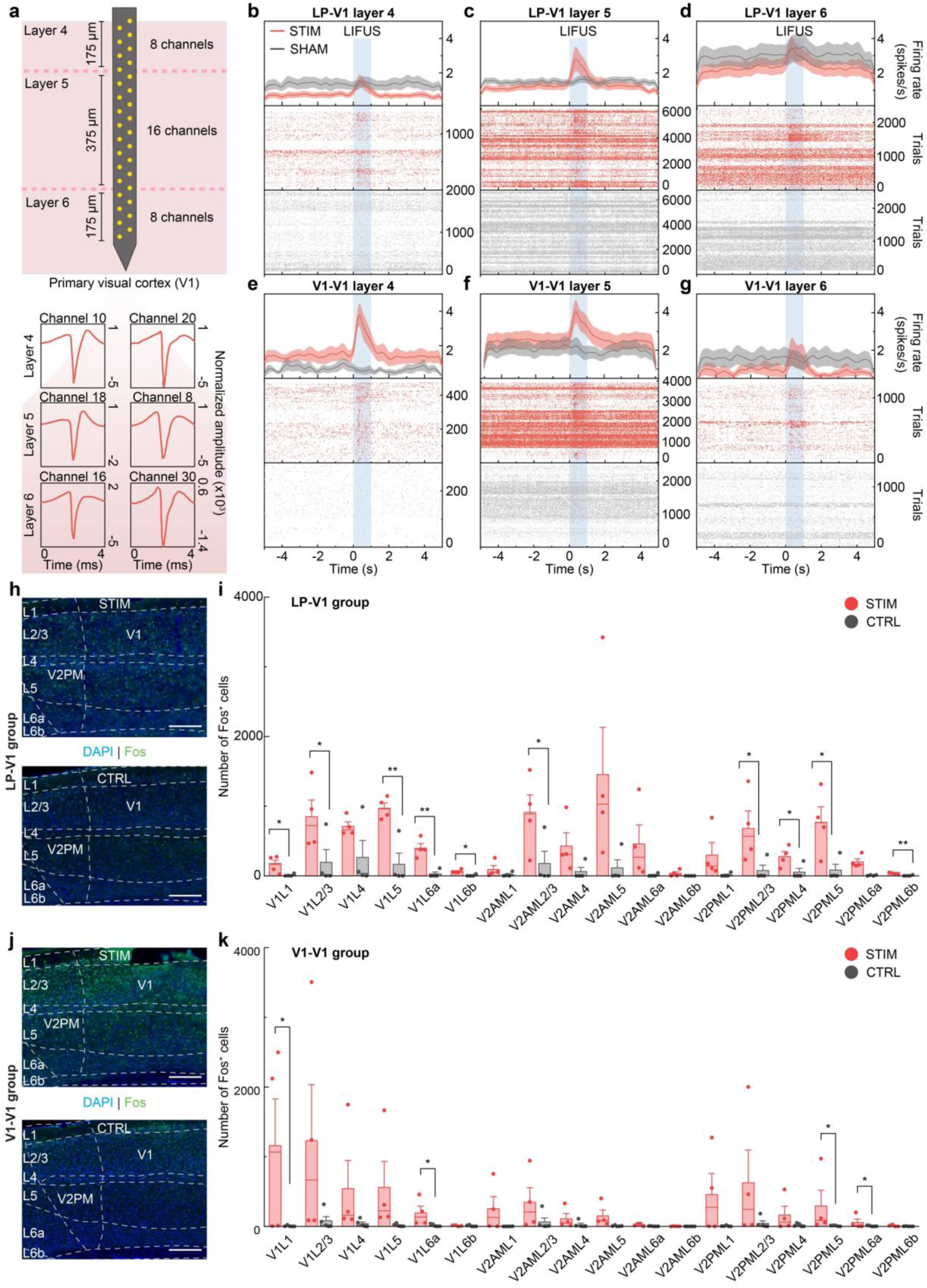
Neural activation across V1 cortical layers. **a**, Schematic indicating the spatial distribution of neural probe electrodes in the V1 (top) and representative spike waveforms in layers 4, 5, and 6 of the V1 (bottom). Single units were recorded from approximately 8 to 16 channels in L4 (DV:-450 to-625 µm), L5 (DV:-625 to-1025 µm), and L6 (DV:-1025 to-1200 µm), covering depths of 175 to 190 µm, 375 to 425 µm, and 175 to 190 µm, respectively. **b-d**, Peri-stimulus time histogram (PSTH, solid line: mean value, shaded area: standard error of the mean (SEM), bin size: 10 ms, Gaussian smoothing SD: 15 ms) and raster plots of single-unit spiking activity in layers 4 (**b**), 5 (**c**), and 6 (**d**) of the V1 for the LP-V1 group STIM and SHAM conditions. Blue shaded bar indicates the 1-s LIFUS duration. **e-g**, Peri-stimulus time histogram (PSTH, solid line: mean value, shaded area: standard error of the mean (SEM), bin size: 10 ms, Gaussian smoothing SD: 15 ms) and raster plots of single-unit spiking activity in layers 4 (**e**), 5 (**f**), and 6 (**g**) of the V1 for the V1-V1 group STIM and SHAM conditions. Blue shaded bar indicates the 1-s LIFUS duration. h, Representative Fos IHC images of the ipsilateral V1 and V2PM layers for LP-V1 group STIM and CTRL conditions. Cortical layers L1 to L6b are demarcated with white dotted lines. Scale bar, 100 µm. **i**, Number of Fos-positive cells in the V1, V2AM, and V2PM cortical layers for the LP-V1 group STIM and CTRL conditions. *p < 0.05, **p < 0.01, two-sided unpaired Student’s t-test, n = 8. **j**, Representative Fos IHC images of the ipsilateral V1 and V2PM layers for V1-V1 group STIM and CTRL conditions. Cortical layers L1 to L6b are demarcated with white dotted lines. Scale bar, 100 µm. **k**, Number of Fos-positive cells in the V1, V2AM, and V2PM cortical layers for the V1-V1 group STIM and CTRL conditions. *p < 0.05, two-sided unpaired Student’s t-test, n = 9.

Our c-Fos IHC results also confirmed this layer-specific circuit modulation using LIFUS of the LP-V1 pathways. Using a custom MATLAB program within our QUINT workflow, we analyzed the Fos-positive cells in cortical layers 1∼ 6 (L1∼ L6) of the V1 and V2 regions of stimulated and control mice (Methods). In the LP-V1 group, significantly higher levels of Fos-positive cells were present in the stimulated mice compared to the control mice in V1L1, V1L2/3, V1L5, V1L6a, V1L6b, V2AML2/3, V2PML2/3, V2PML4, V2PML5, and V2PML6b (Fig. 4h, i and Supplementary Fig. 14a, b; two-tailed unpaired *t*-test, \**p* < 0.05, \*\**p* < 0.01, *n* = 8). We observed the highest Fos-positive cell count in V1L5, V1L6a, and V2PML6b, which is consistent with our electrophysiology results and the LP-V1 neural network. For the V1-V1 group, significantly higher levels of Fos-positive cells were found in the stimulated mice compared to the control mice in V1L1, V1L6a, V2PML5, and V2PML6a (Fig. 4j, k and Supplementary Fig. 14a, b; two-tailed unpaired *t*-test, \**p* < 0.05, *n* = 9), which have been associated with LIFUS-induced cortical activation. These layer-specific responses to LIFUS demonstrate that corticothalamic top-down and bottom-up activation of the LP-V1 circuit are selectively activated by LIFUS and require precise modulation of target regions. This ability to precisely target specific neural circuits is critical for widespread therapeutic applications.

### PRF and region-dependent LIFUS selectively activate E/I neuronal types in the visual system

We next aimed to develop a LIFUS protocol that could selectively modulate excitatory and inhibitory neuronal activity of the LP-V1 pathway. We hypothesized that specific pulse repetition frequencies (PRFs) of ultrasound could have selective effects on different types of neurons and bidirectional circuit pathways. We tested this hypothesis on our LP-V1 and V1-V1 groups by identifying single unit spikes as inhibitory or excitatory (Fig. 5a). The neural spiking units were classified as excitatory (regular spiking units, RSUs) or inhibitory (fast spiking units, FSUs) based on the trough-to-second peak latency of the spike waveforms^60, 61^. We further differentiated the RSUs into RSU1 and RSU2 based on their first-to-second peak ratio (Methods). Consequently, 93, 219, and 175 units were categorized as FSU, RSU1, and RSU2, respectively (Fig. 5b). For the PRF parameters, we designed a waveform with a burst duration of 1 s at an I_SPPA_ of 51.25 W/cm^2^ (peak pressure of 1.24 MPa), 50% duty cycle, and PRFs of 3 kHz, 1 kHz, and 80 Hz (Fig. 5c).

**Fig. 5.**
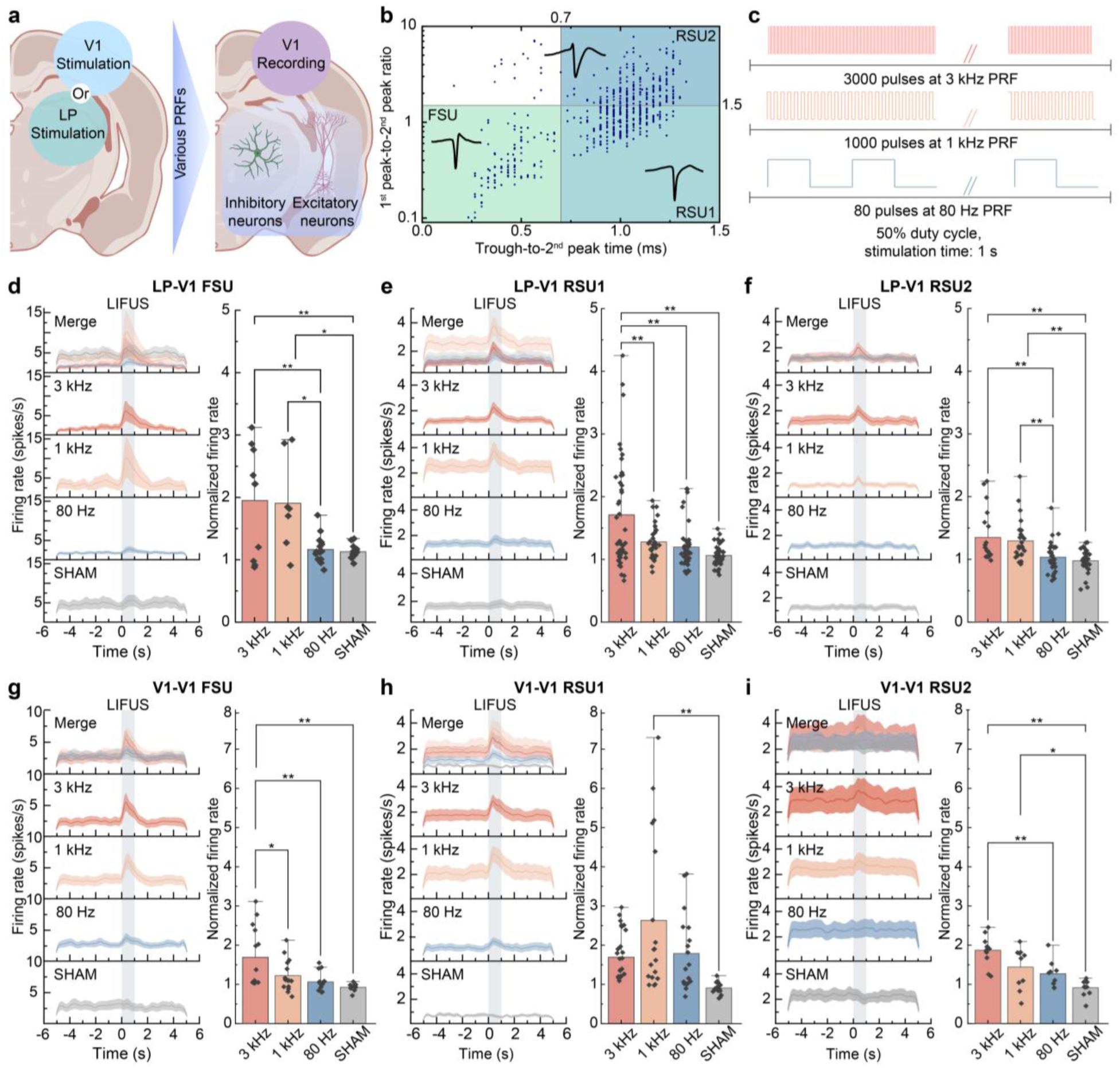
PRF and region-dependent activation of E/I neuronal activity. **a**, Schematic of the PRF and region variation. The LP and V1 were targeted and various PRFs were delivered to selectively activate E/I neuronal pathways in the visual system. **b**, Single-unit classification of 487 units from both the LP-V1 and V1-V1 groups. Each blue dot represents a single unit. Units with a trough-to-second peak latency of less than 0.7 ms were classified as inhibitory fast-spiking units (FSUs; green region with representative waveform). Units with longer latencies were classified as excitatory regular-spiking units (RSUs). RSU1s (light cyan region with representative waveform) and RSU2s (light blue region with representative waveform) were further defined based on their first-to-second peak ratio. **c**, Overview of the ultrasound parameters used in this study. Ultrasound was generated at a PRF of 3 kHz, 1 kHz, and 80 Hz with 3000, 1000, and 80 pulses, respectively, for a 1-s stimulation duration. **d-f**, Peri-stimulus time histograms (PSTHs; solid line: mean value, shaded area: standard error of the mean (SEM), bin size: 10 ms, Gaussian smoothing SD: 15 ms) of neuronal firing rate and bar charts of normalized firing rate of the LP-V1 group. Spike analysis was categorized by FSUs, RSU1s, and RSU2s for 3 kHz, 1 kHz, 80 Hz, and SHAM conditions. The normalized firing rate was compared across the stimulation conditions for the FSU (total of 47 units, *p < 0.05, **p < 0.01, one-way ANOVA with Tukey’s post-hoc test, n = 11) (**d**), RSU1 (total of 173 units, **p < 0.01, one-way ANOVA with Tukey’s post-hoc test, n = 11) (**e**), and RSU2 (total of 112 units, **p < 0.01, one-way ANOVA with Tukey’s post-hoc test, n = 11) (**f**) neuronal types. **g-i**, Peri-stimulus time histograms (PSTHs; solid line: mean value, shaded area: standard error of the mean (SEM), bin size: 10 ms, Gaussian smoothing SD: 15 ms) of neuronal firing rate and bar charts of normalized firing rate of the V1-V1 group. Spike analysis was categorized by FSUs, RSU1s, and RSU2s for 3 kHz, 1 kHz, 80 Hz, and SHAM conditions. The normalized firing rate was compared across the stimulation conditions for the FSU (total 43 units, *p < 0.05, **p < 0.01, one-way ANOVA with Tukey’s post-hoc test, n = 8) (**g**), RSU1 (total 91 units, *p < 0.05, **p < 0.01, one-way ANOVA with Tukey’s post-hoc test, n = 8) (**h**), and RSU2 (total 57 units, *p < 0.05, **p < 0.01, one-way ANOVA with Tukey’s post-hoc test, n = 8) (**j**). Bars represent mean values, and error bars indicate outliers. The grey shaded bar of the PSTH indicates the 1-s stimulation duration.

For the LP-V1 group, FSUs exhibited an increased firing rate in V1 at PRFs of 3 kHz and 1 kHz, which remained elevated throughout the 1-s stimulation compared to the sham condition, while LIFUS at a PRF of 80 Hz did not produce a significant effect (Fig. 5d and Supplementary Fig. 15a). The normalized firing rate during stimulation at 3 kHz and 1 kHz demonstrated a significant rise compared to the 80 Hz and sham conditions, while there was no significant difference in the normalized firing rate during stimulation between the 3 kHz and 1 kHz conditions

(Fig. 5d; one-way ANOVA with Tukey’s *post-hoc* test, *F* = 8.00001, \**p* < 0.05, \*\**p* < 0.01). For RSU1, we also observed a significant increase in the firing rate at PRFs of 3 kHz and 1 kHz compared to the sham condition, while LIFUS at a PRF of 80 Hz did not generate a significant effect (Fig. 5e and Supplementary Fig. 15b). However, for the normalized firing rate, only the 3 kHz PRF showed a significant difference from the 1 kHz, 80 Hz, and sham conditions, indicating that a PRF of 3 kHz was the most effective protocol for activating RSU1 neurons (Fig. 5e; one-way ANOVA with Tukey’s *post-hoc* test, *F* = 13.69416, \*\**p* < 0.01). For RSU2, similar to what was observed in FSU1, PRFs of 3 kHz and 1 kHz induced an increased firing rate (Fig. 5f and Supplementary Fig. 15c) and showed a significant difference in normalized firing rate compared to 80 Hz and sham conditions (Fig. 5f; one-way ANOVA with Tukey’s *post-hoc* test, *F* = 11.24758, \*\**p* < 0.01). In addition, we conducted spectral analysis of the low-pass filtered neural signals at physiologically critical frequency bands of delta (0.1∼ 4 Hz), theta (5∼ 7 Hz), and alpha (8∼ 12 Hz). As expected, the signal power increased during the 1-s LIFUS, particularly for the 3-kHz and 1-kHz PRF conditions across L4, L5, and L6 of the V1 (Supplementary Fig. 16a, b). These results demonstrate that PRFs of 3 kHz and 1 kHz are effective in activating FSU and RSU2 neurons, while 3 kHz is effective for activating RSU1 neurons in bottom-up LP-V1 pathways.

In the V1-V1 group, FSUs in V1 exhibited an increased firing rate during LIFUS at PRFs of 3 kHz, 1 kHz, and 80 Hz, with the increased activity sustained throughout the 1-s stimulation compared to the sham condition (Fig. 5g and Supplementary Fig. 15d). The normalized firing rate for the 3 kHz and 1 kHz conditions demonstrated a significant increase compared to the sham condition. There was no significant difference in the normalized firing rate between the 3 kHz and 1 kHz or between the 80 Hz and sham condition (Fig. 5g; one-way ANOVA with Tukey’s *post-hoc* test, *F* = 12.08249, \**p* < 0.05, \*\**p* < 0.01). For RSU1, we also observed a significant increase in the firing rate for all three PRF conditions compared to the sham condition (Fig. 5h and Supplementary Fig. 15e). However, for the normalized firing rate, only the 1 kHz condition exhibited a significant increase compared to the sham condition, while there was no significant difference between the 3 kHz or 80 Hz and the sham condition, suggesting that a PRF of 1 kHz is the most effective protocol for activating RSU1 (Fig. 5h; one-way ANOVA with Tukey’s *post-hoc* test, *F* = 6.34944, \*\**p* < 0.01). For RSU2, we observed an increase in firing rate for 3 kHz and 1 kHz (Fig. 5i and Supplementary Fig. 15f). This indicated that V1 stimulation at a PRF of 80 Hz activated RSU1 and FSU neurons, but not RSU2 neurons. Analysis of the normalized firing rate revealed that the 3 kHz condition exhibited a significant difference in firing rate compared to the 1 kHz, 80 Hz, and sham conditions, indicating that a PRF of 3 kHz was the most effective at activating RSU2 neurons (Fig. 5i; one-way ANOVA with Tukey’s *post-hoc* test, *F* = 7.0426, \**p* < 0.05, \*\**p* < 0.01). We also conducted spectral analysis for the V1-V1 group and found that the signal power increased during the 1-s LIFUS, particularly for the 3-kHz and 1-kHz PRF conditions across L4, L5, and L6 of the V1 (Supplementary Fig. 16a, b). Interestingly, the 1-kHz PRF condition elicited a stronger response in the delta and theta bands compared to the 3-kHz PRF condition, suggesting a PRF-dependent mechanism for modulating low-frequency brain oscillations. These results demonstrate that PRFs of 3 kHz and 1 kHz are effective in activating FSU neurons, while 1 kHz is effective for activating RSU1 neurons and 3 kHz is most effective at activating RSU2 neurons in top-down LP-V1 pathways.

Comparing the LP-V1 and V1-V1 groups, we found that for activating FSUs in V1, PRF of 3 kHz and 1 kHz were shown to be effective protocols when stimulating either the LP or V1 regions. Similarly, for activating RSU2s in the V1, PRF of 3 kHz and 1 kHz were effective protocols when stimulating the LP, while only the 3 kHz condition was effective when stimulating the V1. In addition, we observed that 3 kHz was the most effective protocol for activating RSU1s in V1 when stimulating the LP, whereas 1 kHz was the most effective for activating RSU1s in V1 when stimulating the V1. These electrophysiological results suggest that modulation of the E/I balance in the LP-V1 circuit is more effective using higher PRF LIFUS, and that those responses are also governed by top-down and bottom-up region-dependent corticothalamic pathways. In particular, bottom-up activation of excitatory RSU and inhibitory FSU was most effective by targeting the thalamus at a PRF of 3 kHz.

### LIFUS can modulate synaptic transmission via specific neurotransmitter release in bidirectional pathways of the LP-V1 circuit

While FSUs and RSUs provide crucial information on the functional roles of neural circuits such as network synchronization and integration of signal input, they lack biochemical characterizations such as synaptic signaling, which broadly influences brain states and behaviors^28, 62, 63^. Thus, based on our electrophysiological results, we next explored which specific neurotransmitter was associated with the FSU and RSU responses. We employed an immunohistochemistry (IHC)-based approach to tag two main neurotransmitters in the brain, vGLUT1 and GABA, which are responsible for excitatory and inhibitory neuronal activity, respectively. As the neurotransmitter for glutamatergic neurons, vGLUT1 is one of the primary synaptic proteins essential for excitatory neurotransmission in the central nervous system. vGLUT1 is ubiquitously found in the central nervous system-predominantly in the cortex, hippocampus, and thalamus-and is often implicated in higher-order cognitive functions^63^. GABA is the primary inhibitory neurotransmitter in the brain and is found in the soma as well as synaptic gaps of neurons. The presence of GABA generally indicates GABAergic inhibitory interneurons. To determine LIFUS-induced activation of neurotransmitters and corresponding neuronal types, we co-stained the LP-V1 and V1-V1 groups using Fos-GABA and Fos-vGLUT1 antibody pairs for control, sham, 3 kHz PRF, 1 kHz PRF, and 80 Hz PRF groups.

For the LP-V1 group, we found that LIFUS of LP delivered at a PRF of 3 kHz significantly increased glutamatergic activity in the V1, while GABAergic activity remained unchanged compared to control and sham levels across all stimulation conditions (Fig. 6a-c and Supplementary Fig. 17a, b; one-way ANOVA with Tukey’s *post-hoc* test, *F* = 4.89142, \**p* < 0.05). Similarly, LP stimulation at a PRF of 3 kHz significantly induced glutamatergic activity in the LP, while GABAergic activity was nonspecific to all PRF parameters (Fig. 6d, e and Supplementary Fig. 17a, b; one-way ANOVA with Tukey’s *post-hoc* test, *F* = 5.67807, \*\**p* < 0.01, *n* = 5). This indicates that LP stimulation can selectively activate glutamatergic neurons in both the LP and V1 regions. On the other hand, for the V1-V1 group, LIFUS of V1 triggered at a PRF of 3 kHz or 1 kHz significantly increased glutamatergic activity in the V1 region, while GABAergic activity was not significantly affected by any PRF parameter (Fig. 6f-h and Supplementary Fig. 18a, b; one-way ANOVA with Tukey’s *post-hoc* test, *F* = 13.68501, \**p* < 0.05, *n* = 5). Interestingly, the 3 kHz condition activated a significantly higher level of glutamatergic neurons compared to the 1 kHz condition, indicating that neuronal activation due to direct cortical stimulation can be controlled with PRFs in addition to intensity (Fig. 6h; one-way ANOVA with Tukey’s *post-hoc* test, *F* = 13.68501, \**p* < 0.05). In the LP, LIFUS of V1 did not induce any significant GABAergic or glutamatergic activity compared to the control and sham levels for all stimulation conditions (Fig. 6i, j and Supplementary Fig. 18a, b). This suggests that top-down modulation of the LP-V1 circuit using LIFUS is limited in neuronal specificity.

**Fig. 6.**
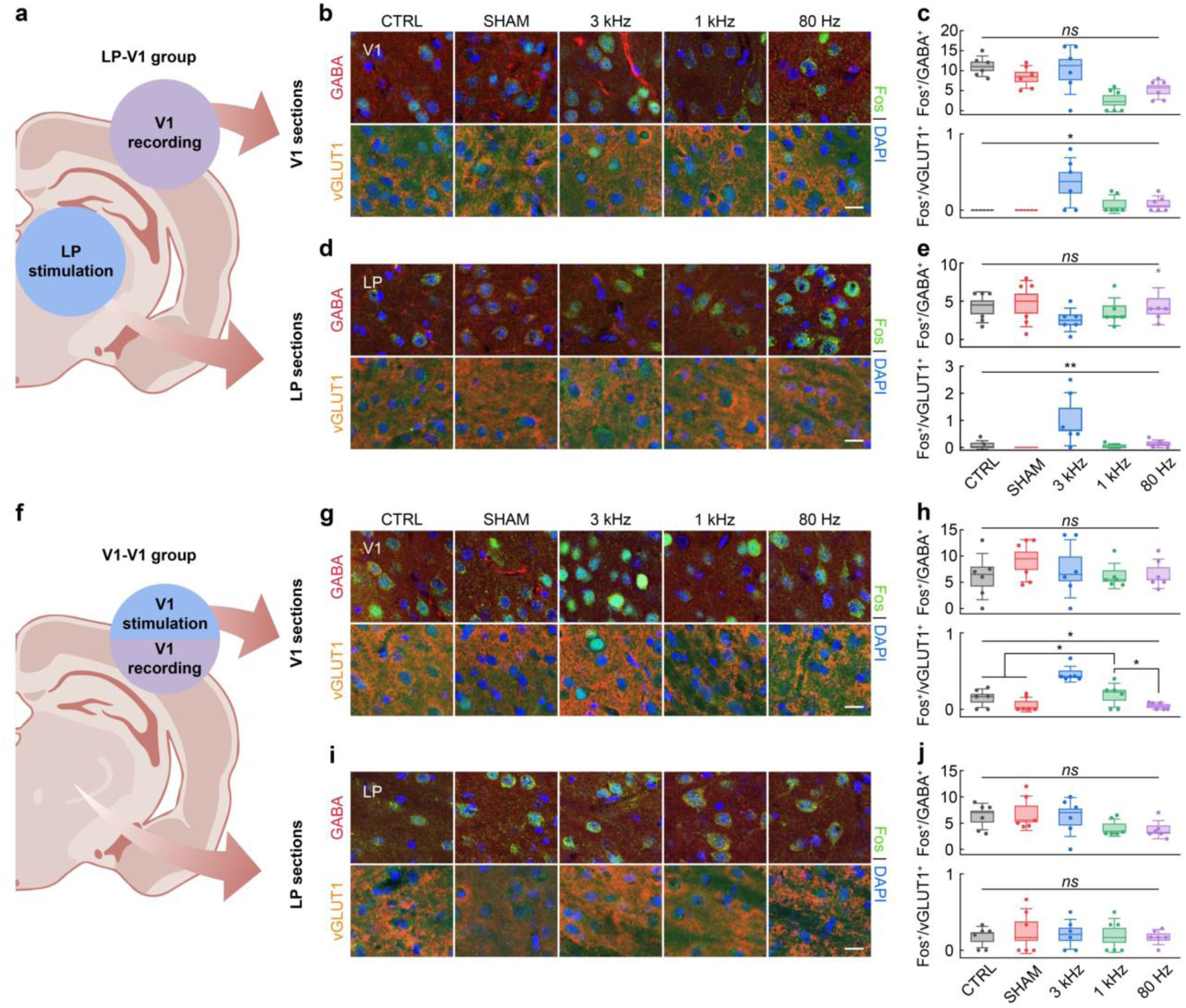
PRF-dependent activation of glutamatergic and GABAergic neurons. **a**, Schematic of the LP-V1 group. **b**, LP-V1 group: Representative IHC images of GABA and vGLUT1 co-stained with Fos in the V1 for CTRL, SHAM, 3 kHz, 1 kHz, and 80 Hz conditions. Scale bar, 10 µm. **c**, LP-V1 group: Number of Fos-positive GABAergic and glutamatergic cells in the V1 for CTRL, SHAM, 3 kHz, 1 kHz, and 80 Hz conditions. ns, not statistically significant. *p < 0.05, one-way ANOVA with Tukey’s post-hoc test, n = 6. **d**, LP-V1 group: Representative IHC images of GABA and vGLUT1 co-stained with Fos in the LP for CTRL, SHAM, 3 kHz, 1 kHz, and 80 Hz conditions. Scale bar, 10 µm. **e**, LP-V1 group: Number of Fos-positive GABAergic and glutamatergic cells in the LP for CTRL, SHAM, 3 kHz, 1 kHz, and 80 Hz conditions. ns, not statistically significant. **p < 0.01, one-way ANOVA with Tukey’s post-hoc test, n = 6. **f**, Schematic of the V1-V1 group. **g**, V1-V1 group: Representative IHC images of GABA and vGLUT1 co-stained with Fos in the V1 for CTRL, SHAM, 3 kHz, 1 kHz, and 80 Hz conditions. Scale bar, 10 µm. **h**, V1-V1 group: Number of Fos-positive GABAergic and glutamatergic cells in the V1 for CTRL, SHAM, 3 kHz, 1 kHz, and 80 Hz conditions. ns, not statistically significant. *p < 0.05, one-way ANOVA with Tukey’s post-hoc test, n = 6. **i**, V1-V1 group: Representative IHC images of GABA and vGLUT1 co-stained with Fos in the LP for CTRL, SHAM, 3 kHz, 1 kHz, and 80 Hz conditions. Scale bar, 10 µm. **j**, V1-V1 group: Number of Fos-positive GABAergic and glutamatergic cells in the LP for CTRL, SHAM, 3 kHz, 1 kHz, and 80 Hz conditions. ns, not statistically significant, one-way ANOVA with Tukey’s post-hoc test, n = 6.

These findings provide experimental evidence that LIFUS delivered at a PRF of 3 kHz can preferentially modulate bottom-up glutamatergic pathways of the corticothalamic visual circuit, and brain region selection can modulate glutamatergic pathways of the corticocortical visual circuit. V1-V1 top-down LIFUS also shows the potential of modulating the strength of glutamatergic neuronal activity with PRF in addition to acoustic intensity. Furthermore, our results demonstrate that the FSU responses observed in the electrophysiological data are not directly related to GABAergic activation, while the RSU responses strongly suggest a glutamatergic role. This understanding of both electrophysiological and synaptic neurotransmission characteristics of LIFUS is crucial in developing a precise stimulation protocol for circuit-specific modulation.

## Discussion

In this work, we demonstrated selective neuromodulation of E/I neurons in the top-down and bottom-up corticothalamic pathways of the visual system using LIFUS. Based on previous studies, we hypothesized that region and waveform-dependent parameters were crucial for precise neuromodulation. We investigated the electrophysiological response of FSU and RSU neurons, as well as the synaptic neurotransmission of GABA and vGLUT1 to LIFUS delivered at various PRFs. Our results confirmed that PRF and region specificity of LIFUS play major roles in neuron-specific and bidirectional circuit activation.

Stimulation of the LP (LP-V1 group) or V1 (V1-V1 group) induced an acute neurological response in the V1 as measured by an increase in the neuronal spike rate during ultrasound delivery. In both the LP-V1 and V1-V1 groups, LIFUS selectively activated the LP-V1 circuit as verified by brain-wide Fos-positive cells in the LP, dLGN, vLGN, V1, V2AM, V2PM, and SC regions. For LP stimulation, a PRF of 3 kHz and 1 kHz was most effective at activating FSU and RSU2 neurons in the V1, while 3 kHz was effective for activating RSU1 neurons in the V1. On the other hand, for the V1 simulation, a PRF of 3 kHz was most effective at activating FSU and RSU2 neurons in the V1, while 1 kHz was effective at activating RSU1 neurons in the V1. Furthermore, using IHC methods to co-stain for GABA and vGLUT1 neurotransmitters with c-Fos, we discovered that a PRF of 3 kHz selectively activated glutamatergic neurons in the LP and V1 when targeting the LP. When targeting the V1, however, there was a negligible difference of GABAergic and glutamatergic neuronal activation levels across PRF conditions in the LP. In the V1, a PRF of 3 kHz provided the most effective glutamatergic neuronal activation, which is in line with previous studies of cortical stimulation using ultrasound^26^. In general, for visual corticothalamic activation, our findings suggest that targeting the thalamus instead of the visual cortex provides more effective E/I modulation in bidirectional visual pathways.

These results indicate that PRF and region-specificity may play a greater role in selective activation of neuronal types, response dynamics, and synaptic signaling than previously reported. Top-down and bottom-up E/I pathways respond effectively to select PRFs depending on the target region, which suggests that LIFUS requires brain region-optimized parameters in addition to neuron type-optimized parameters for precise circuit activation. This is further corroborated by our IHC data, which demonstrates preferential activation of bottom-up glutamatergic neurons at 3 kHz. Extensive PRF screening and protein marker analysis will be required for a comprehensive map of functional LIFUS activation of the brain. In particular, additional neurotransmitters such as vGLUT2 and VGAT (vesicular GABA transporter) could significantly enhance the localization of glutamatergic and GABAergic neurons, respectively.

The functional selectivity of LIFUS to neuronal type and corresponding pathway-dependent neuromodulation is critical for behavioral and therapeutic studies, which rely on circuit-specific physiology. However, while our results demonstrate the critical role of PRF for circuit-specific neuromodulation in the visual system, the underlying molecular mechanism remains unclear. Furthermore, the confounding effects and relationship between anesthesia level and LIFUS intensity require additional investigations. Anesthesia type and stability have been shown to affect LIFUS-induced responses in electrophysiological and behavioral readouts^64–67^. This is particularly critical for circuit-specific applications that require brain-wide connectivity. Nonetheless, this work demonstrates a crucial observation of selective GABAergic and glutamatergic neuronal activation and E/I single unit activity in a specific neural circuit for a wide range of PRFs *in vivo*. Future works could apply this method to precisely manipulate vision-specific behaviors across a wide spectrum of brain disorders, which could accelerate the development of precision therapeutic protocols for circuit-specific diseases.

## Acknowledgements

This research was supported by the K-Brain Project of the National Research Foundation (NRF) funded by the Korean government (MSIT) (RS-2023-00262568), by a grant of the Korea Dementia Research Project through the Korea Dementia Research Center (KDRC), funded by the Ministry of Health & Welfare and Ministry of Science and ICT, Republic of Korea (RS-2024-00355871), by National R&D Program through the National Research Foundation of Korea (NRF) funded by Ministry of Science and ICT (2020M3H2A107804521), by KBRI basic research program through Korea Brain Research Institute funded by Ministry of Science and ICT (24-BR-05-02), and by BK21 FOUR (Connected AI Education & Research Program for Industry and Society Innovation, KAIST EE, No. 4120200113769). We thank Dr. Maesoon Im (KIST) for his initial role in choosing a brain circuit to target. We also thank Prof. Byungkook Lim (UCSD) and Prof. Seung-Hee Lee (KAIST) for their advice and guidance in interpreting our biological results. In addition, we would like to acknowledge the facilities, and the scientific and technical assistance of the EM & Histology Core Facility and Dr. Yongsuk Hur at the BioMedical Research Center, KAIST.

## CRediT authorship statement

**Yehhyun Jo:** Conceptualization, Methodology, Software, Validation, Formal analysis, Investigation, Resources, Data Curation, Writing - Original Draft, Writing - Review & Editing, Visualization. **Xiaojia Liang:** Conceptualization, Methodology, Software, Validation, Formal analysis, Investigation, Data Curation, Writing - Original Draft, Writing - Review & Editing, Visualization. **Hong Hanh Nguyen:** Software, Validation, Formal analysis, Data Curation. **Yeonseo Choi:** Validation, Formal analysis, Investigation. **Ga-Eun Bae:** Validation, Investigation. **Yakdol Cho:** Resources. **Jiwan Woo:** Resources. **Hyunjoo Jenny Lee:** Conceptualization, Methodology, Formal analysis, Resources, Writing - Original Draft, Writing - Review & Editing, Supervision, Project administration, Funding acquisition.

