## Supplementary Information for "Selective manipulation of excitatory and inhibitory neurons in top-down and bottom-up visual pathways using ultrasound stimulation"

**†These authors contributed equally**


### Supplementary Figures 1~ 18

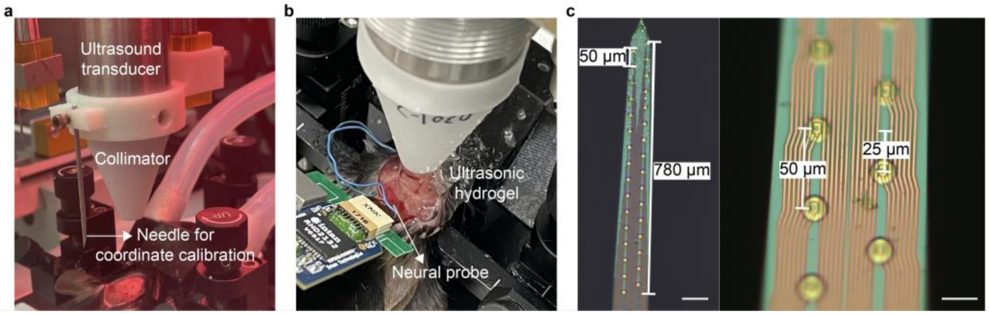


*Supplementary Fig. 1 | In vivo experimental setup.* ***a****, Ultrasound transducer with a collimator and calibration needle for precise targeting of LIFUS.* ***b****, Ultrasound transducer with collimator positioned vertically over the target region with a silicon neural probe inserted into the visual cortex at a 40-degree angle from the sagittal plane.* ***c****, Optical microscope images of the 32-channel silicon neural probe. Scale bar, 70 µm (left), 25 µm (right).*

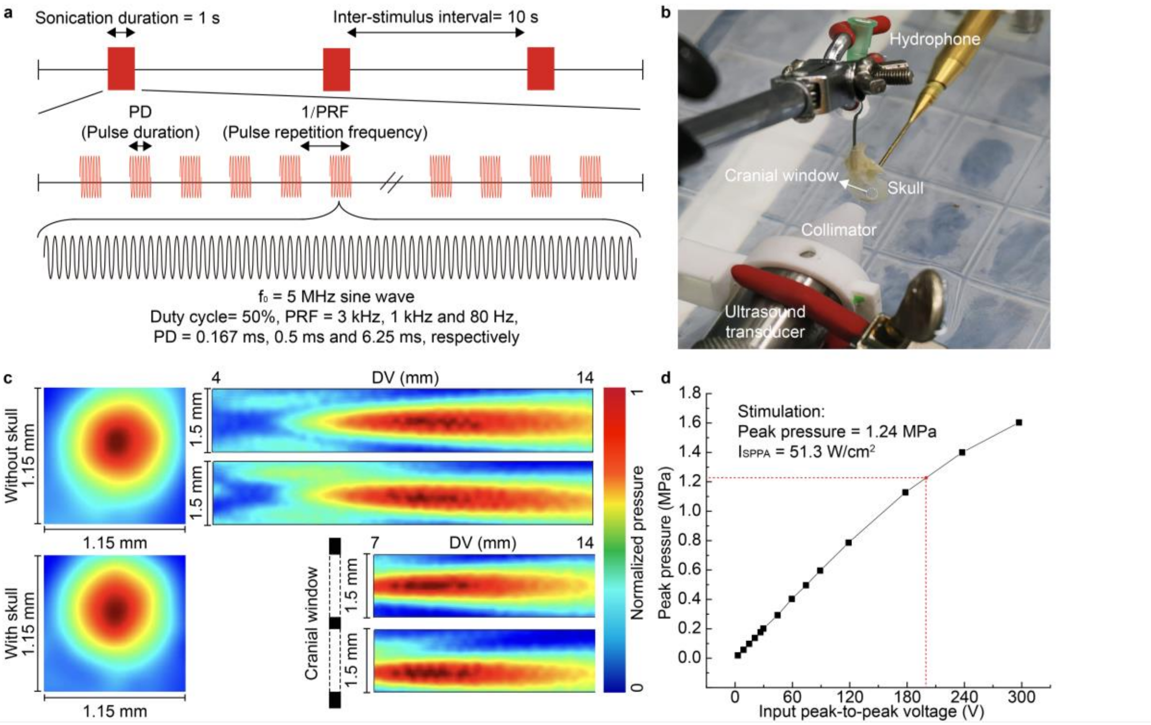


*Supplementary Fig. 2 | Ultrasound stimulation protocol and beam profile.* ***a***, *Schematic of the ultrasound waveform used for stimulation.* ***b***, *Photograph of the beam profile measurement setup with an ex vivo mouse skull.* ***c***, *Beam profile measurements without the skull and with a craniotomized skull. The color bar represents normalized pressure.* ***d***, *Saturation curve of the peak output pressure of the transducer with increasing input voltage. The peak pressure at a driving voltage of 204 V was 1.24 MPa.*

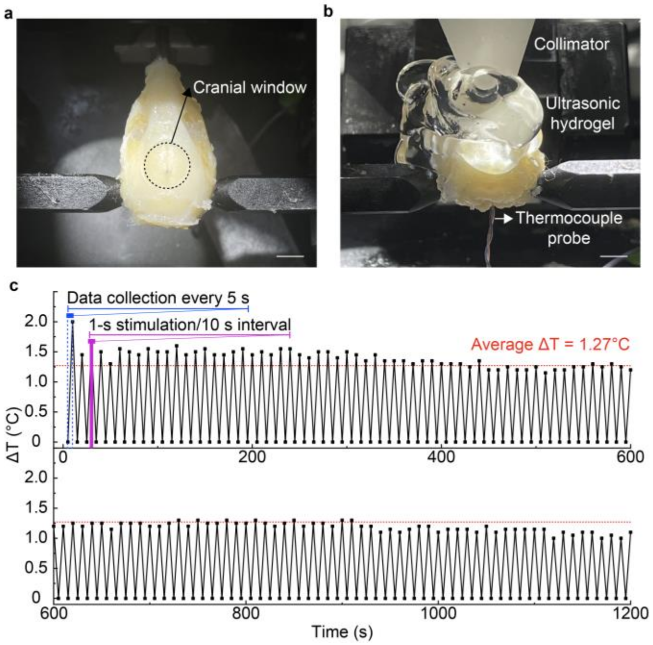


*Supplementary Fig. 3 | Thermocouple test. **a**, Ex vivo brain phantom with perfused brain and craniotomized skull. Scale bar, 3.8 mm. **b**, Experimental setup for monitoring temperature change within the brain phantom during LIFUS using a thermocouple. Scale bar, 3mm. **c**, Plot of temperature change in the brain over a 20-min period of 1-s LIFUS delivered every 10 s.*

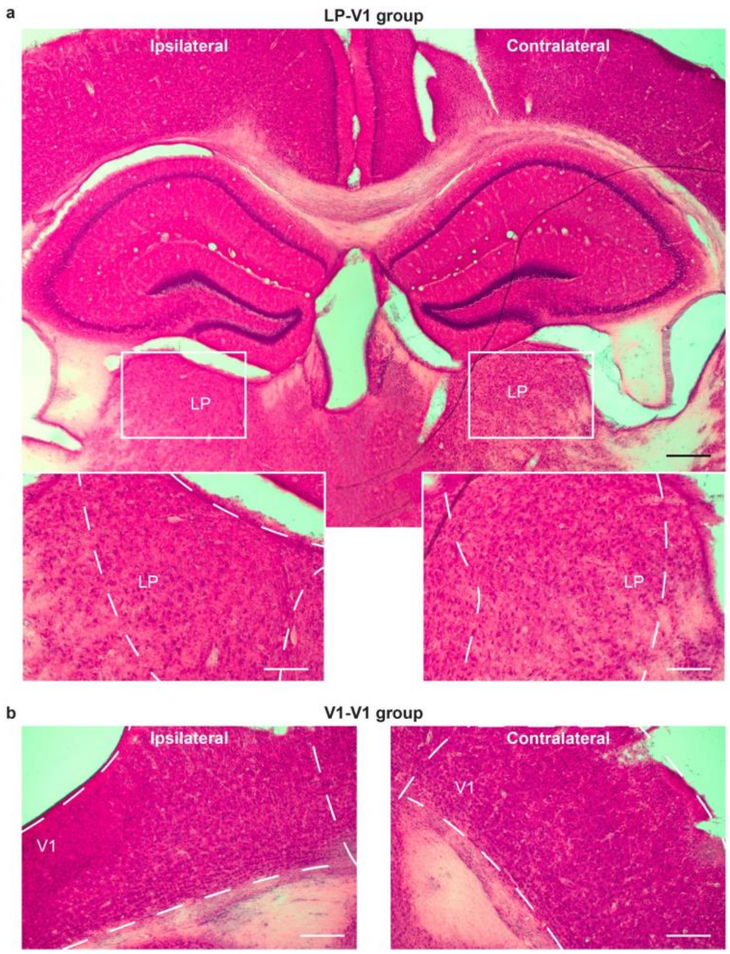


*Supplementary Fig. 4 | Brightfield images of H&E-stained brain sections.* ***a****, Representative H&E images of the LP region for the LP-V1 group. Scale bar, 400 µm.* ***b****, Representative H&E images of the V1 region for the V1-V1 group. Scale bar, 400 µm.*

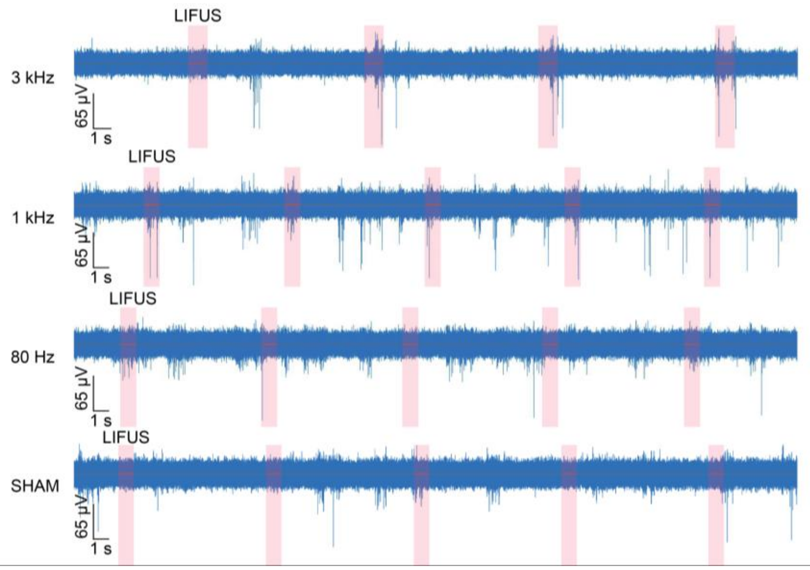


*Supplementary Fig. 5 | Representative raw signals of multi-unit activity in the V1 during LIFUS. Red shaded bar indicates 1-s stimulation duration.*

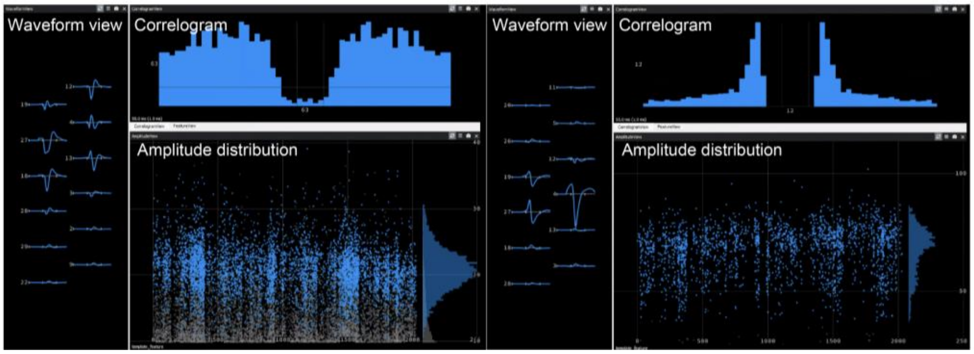


*Supplementary Fig. 6 | Representative images of spike-sorted single units in Phy after running analysis with Kilosort 2.5.*

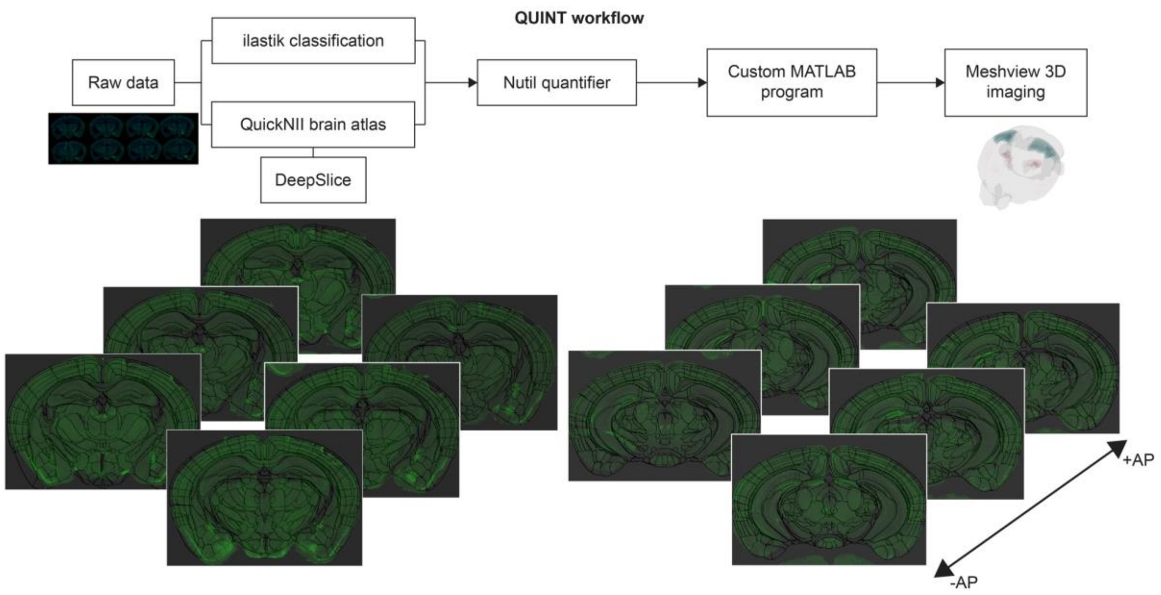


*Supplementary Fig. 7 | Schematic of the QUINT workflow for brain-wide Fos analysis. The pipeline includes ilastik, QuickNII, DeepSlice, Nutil, and MATLAB.*

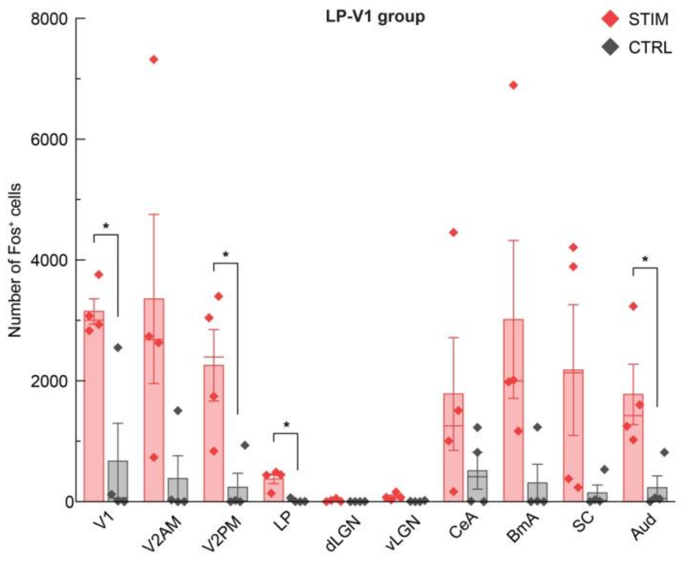


*Supplementary Fig. 8 | Number of Fos-positive cells in the V1, V2AM, V2PM, LP, dLGN, vLGN, CeA, BmA, SC, and Aud regions for the LP-V1 group STIM and CTRL conditions. *p < 0.05, two-sided unpaired Student's t-test, n = 8.*

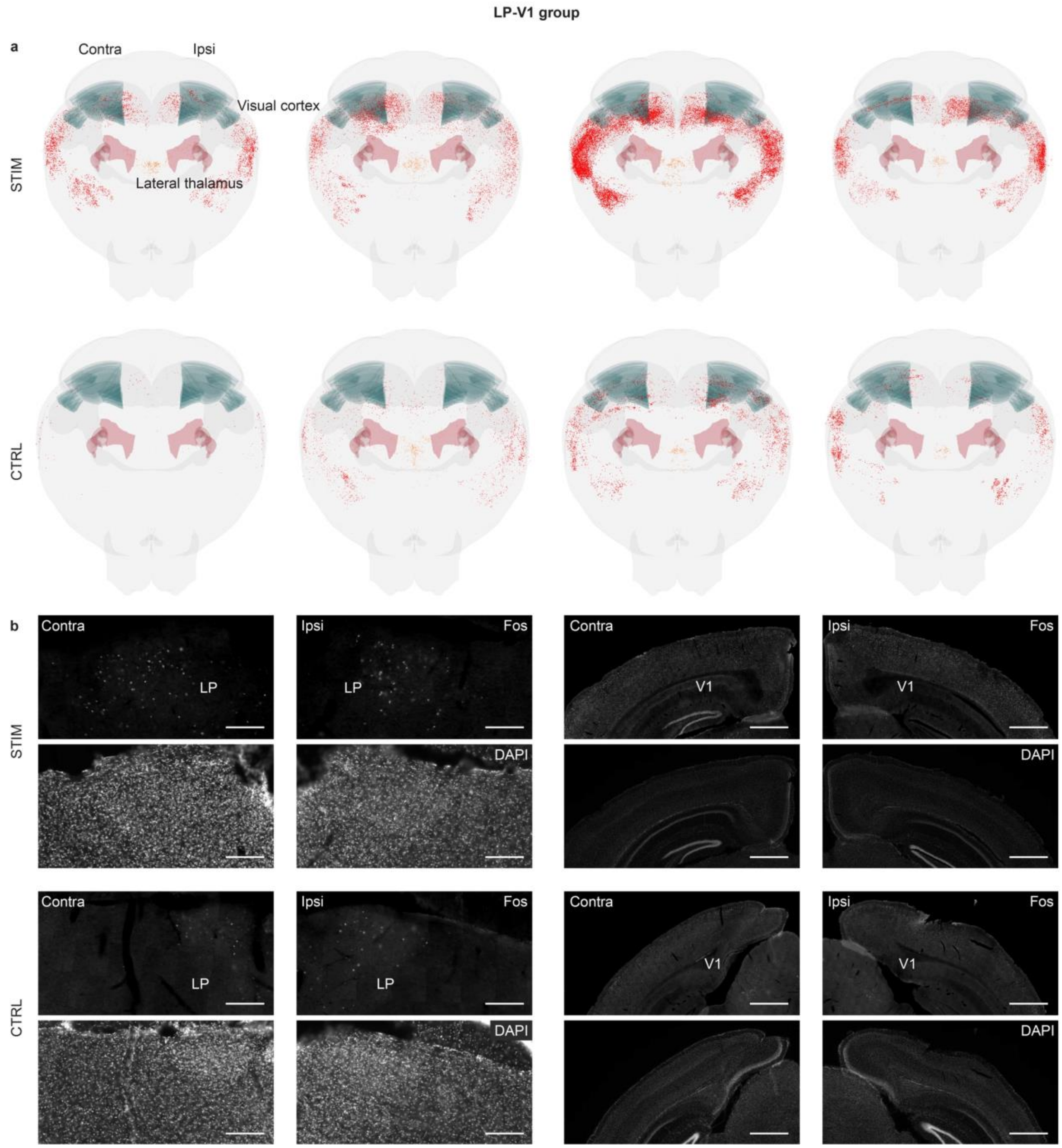


*Supplementary Fig. 9 | Fos IHC analysis for the LP-V1 group.* ***a***, *Representative 3D mesh views of Fos-positive cells in the thalamus and visual cortex for STIM and CTRL conditions.* ***b***, *Representative Fos IHC images of the ipsilateral and contralateral LP and V1 regions for STIM and CTRL conditions. Scale bar (left), 150 µm. Scale bar (right), 800 µm.*

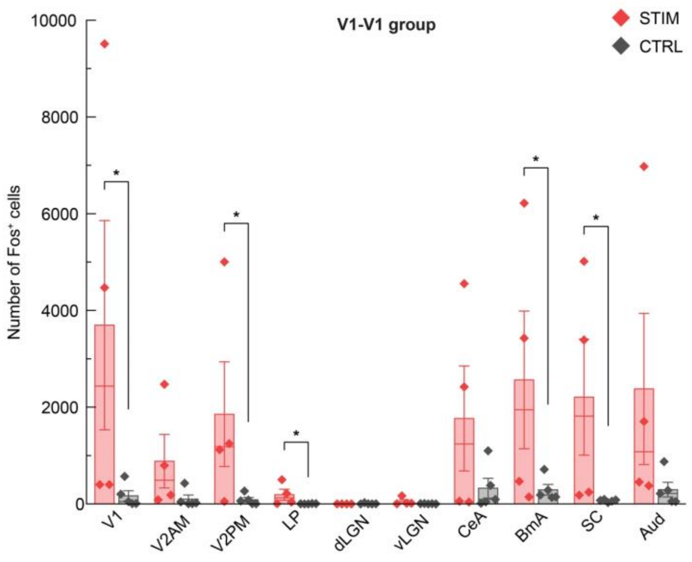


*Supplementary Fig. 10 | Number of Fos-positive cells in the V1, V2AM, V2PM, LP, dLGN, vLGN, CeA, BmA, SC, and Aud regions for the V1-V1 group STIM and CTRL conditions. *p < 0.05, two-sided unpaired Student's t-test, n = 8.*

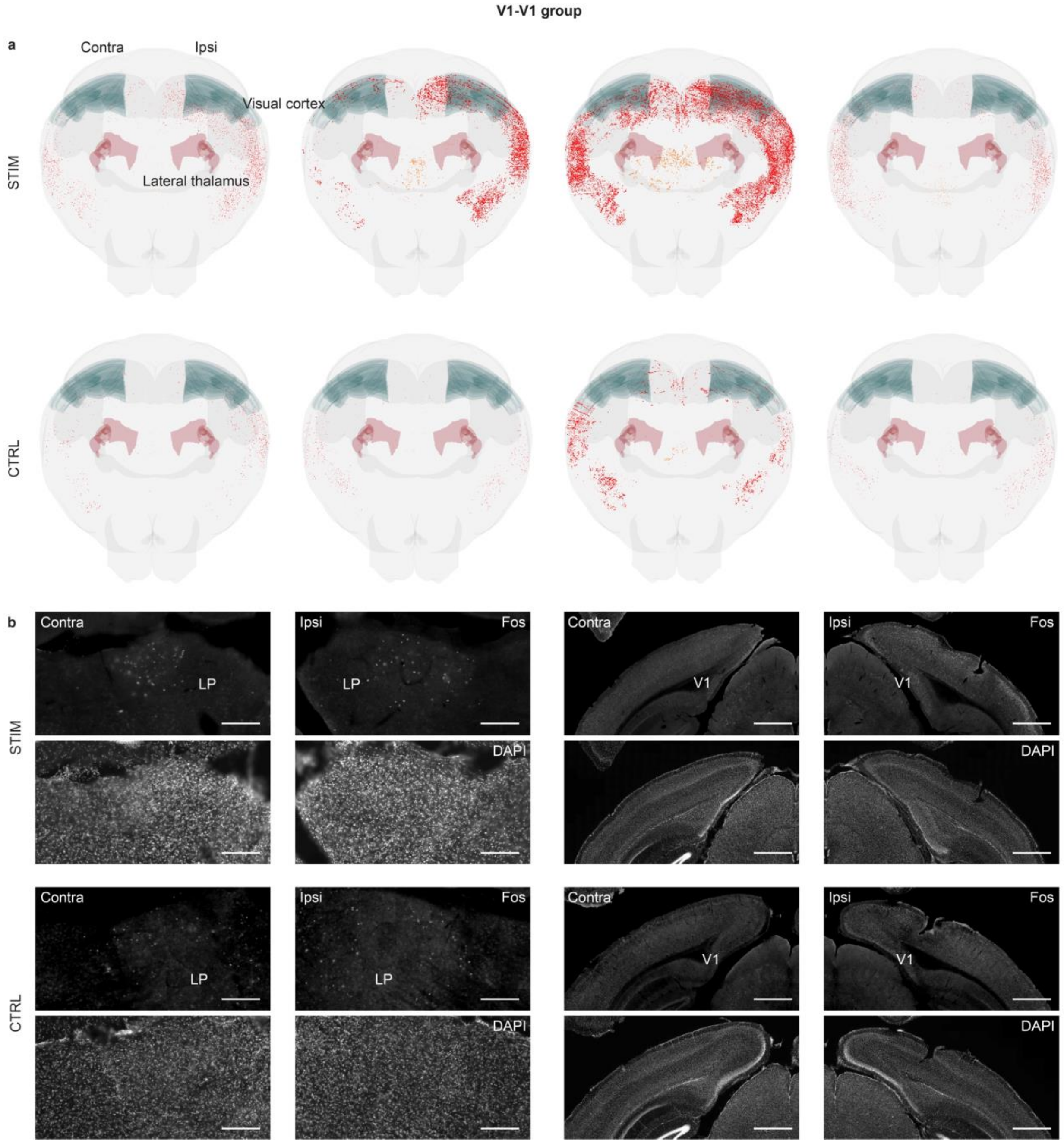


*Supplementary Fig. 11 | Fos IHC analysis for the V1-V1 group.* ***a****, Representative 3D mesh views of Fos-positive cells in the thalamus and visual cortex for STIM and CTRL conditions.* ***b****, Representative Fos IHC images of the ipsilateral and contralateral LP and V1 regions for STIM and CTRL conditions. Scale bar (left), 150 µm. Scale bar (right), 800 µm.*

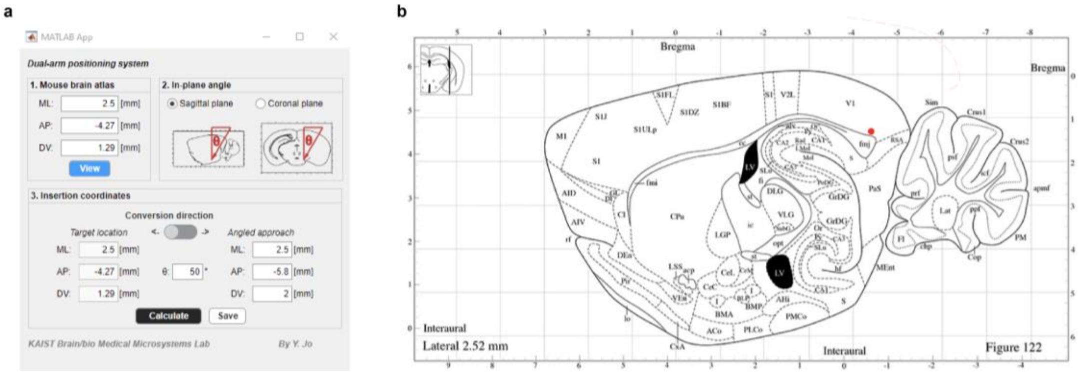


*Supplementary Fig. 12 | Custom MATLAB GUI for calculating the insertion position of a neural probe.* ***a****, Design of the custom GUI in MATLAB, along with an example of the positioning result generated after execution.* ***b****, Representative image indicating the position of the neural probe in the brain, aligned with a version of the Allen Brain Atlas provided by Matt Gaidica. The red dot indicates the tip of the neural probe.*

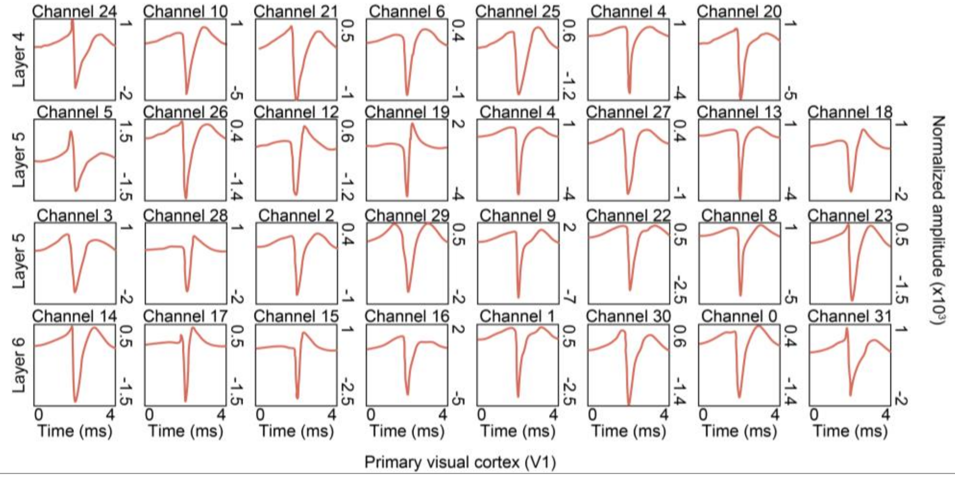


*Supplementary Fig. 13 | Representative spike waveforms of each channel of the neural probe recorded from the visual cortex. Waveform amplitudes are normalized values obtained after analysis with Kilosort 2.5.*

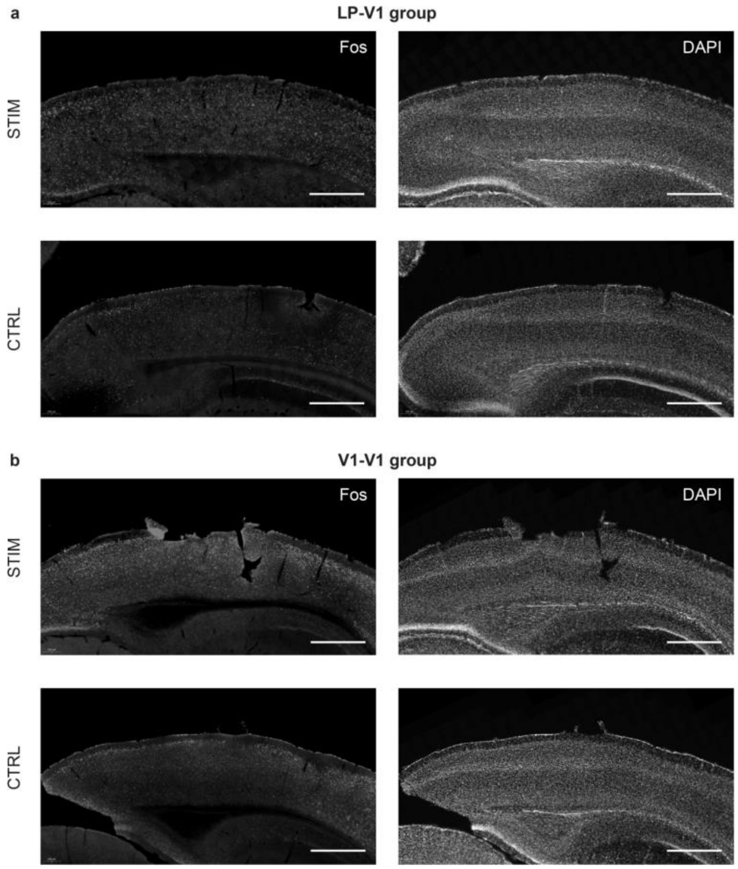


*Supplementary Fig. 14 | Fos IHC analysis of the visual cortical layers.* ***a****, Representative Fos IHC images of the ipsilateral visual cortical layers for LP-V1 group STIM and CTRL conditions. Scale bar, 800 µm.* ***b****, Representative Fos IHC images of the ipsilateral visual cortical layers for V1-V1 group STIM and CTRL conditions. Scale bar, 800 µm.*

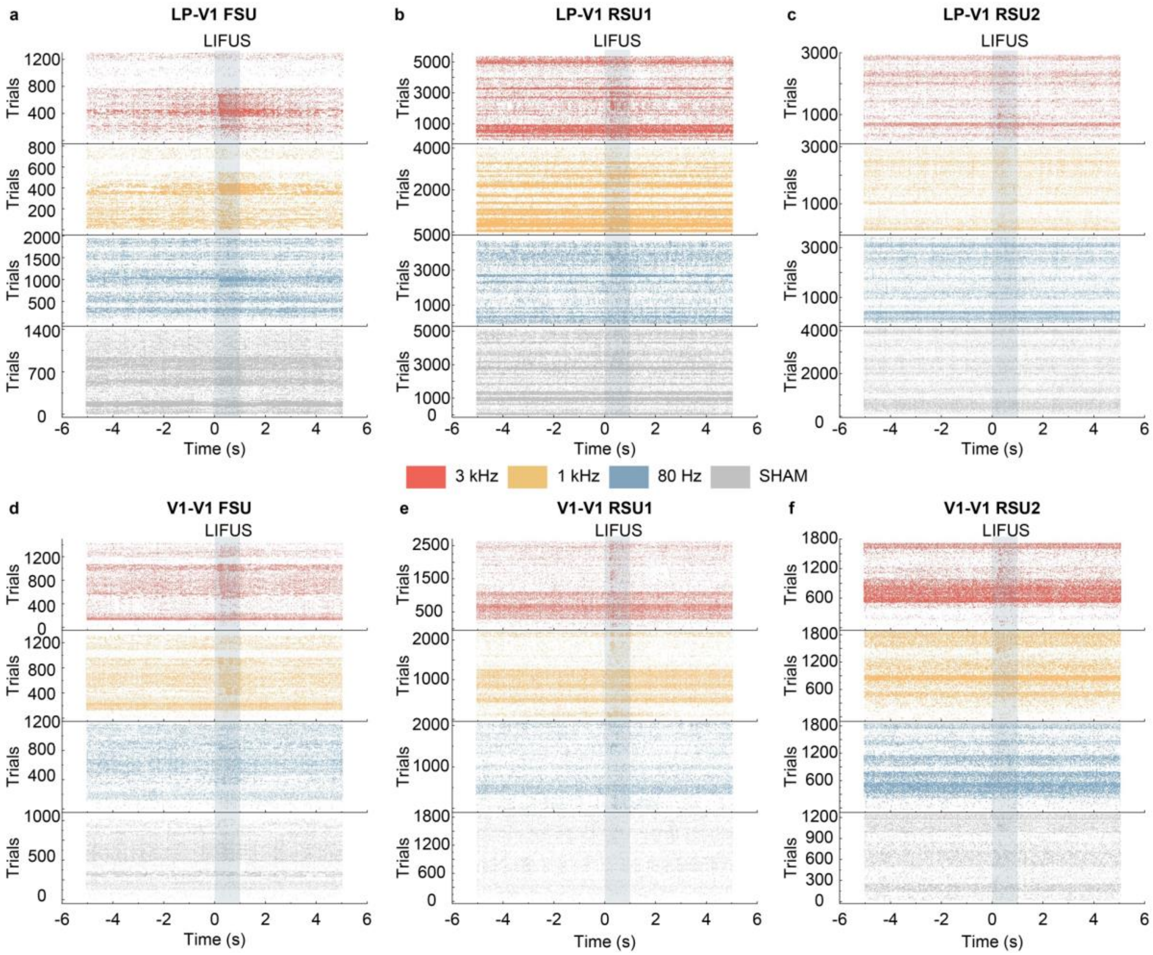


*Supplementary Fig. 15 | Raster plots of neural activity at various PRFs for the LP-V1 and V1-V1 groups.* ***a-c****, Raster plots for the LP-V1 group. Spike analysis was categorized by FSUs (****a****), RSU1s (****b****), and RSU2s (****c****) for 3 kHz, 1 kHz, 80 Hz, and SHAM conditions.* ***d-f****, Raster plots for the V1-V1 group. Spike analysis was categorized by FSUs (****d****), RSU1s (****e****), and RSU2s (****f****) for 3 kHz, 1 kHz, 80 Hz, and SHAM conditions. Grey shaded bar indicates the 1-s LIFUS duration.*

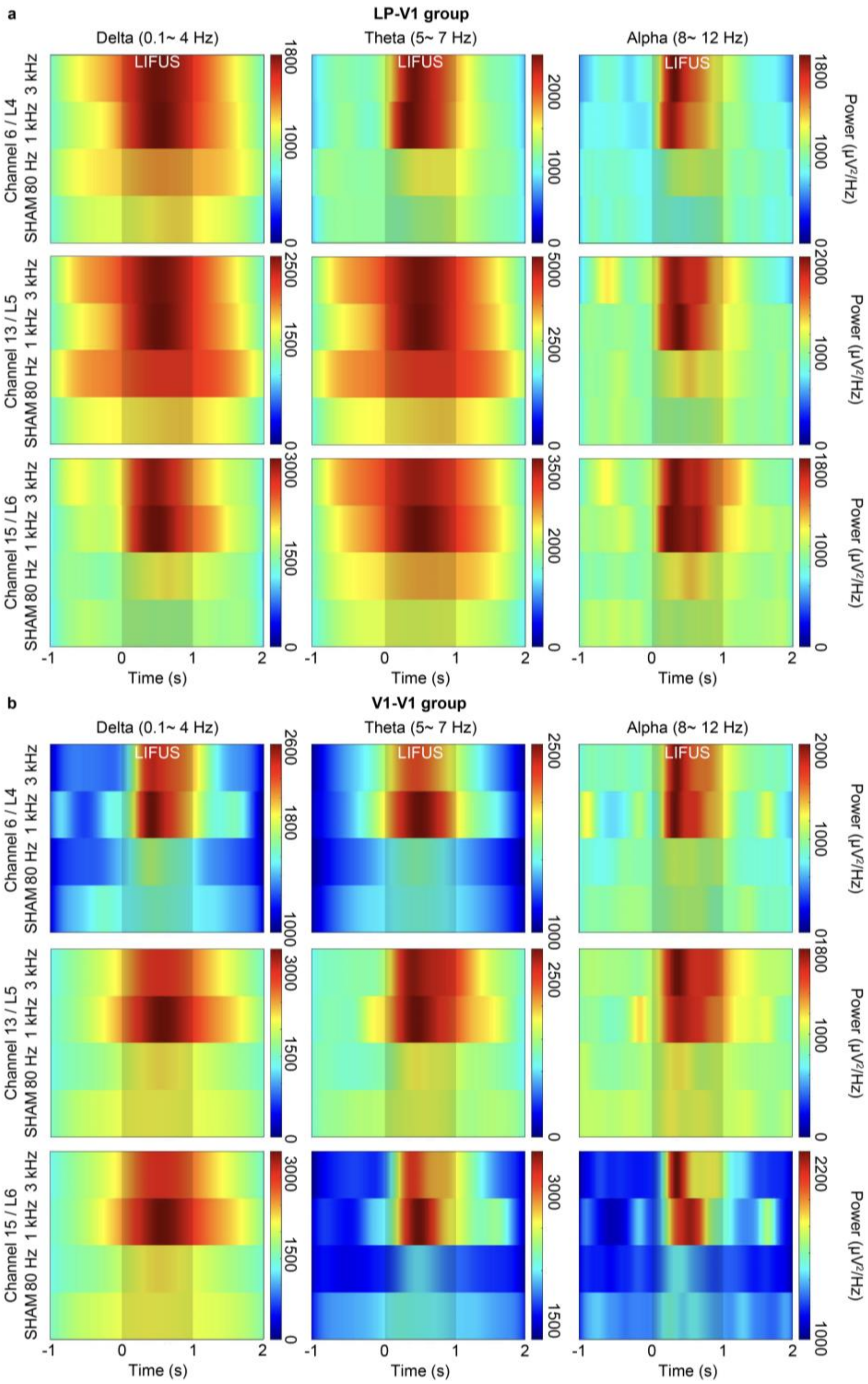


*Supplementary Fig. 16 | Spectrogram of LFP signals across various PRFs.* ***a***, *Spectrograms of delta, theta, and alpha frequency bands for representative neural probe channels located in layer 4, layer 5, and layer 6 of the V1 for the LP-V1 group. The power spectrum during LIFUS is compared across 3 kHz, 1 kHz, 80 Hz, and SHAM conditions.* ***b***, *Spectrograms of delta, theta, and alpha frequency bands for representative neural probe channels located in layer 4, layer 5, and layer 6 of the V1 for the V1-V1 group. The power spectrum during LIFUS is compared across 3 kHz, 1 kHz, 80 Hz, and SHAM conditions. The shaded area between time 0 and 1 s indicates the 1-s stimulation duration.*

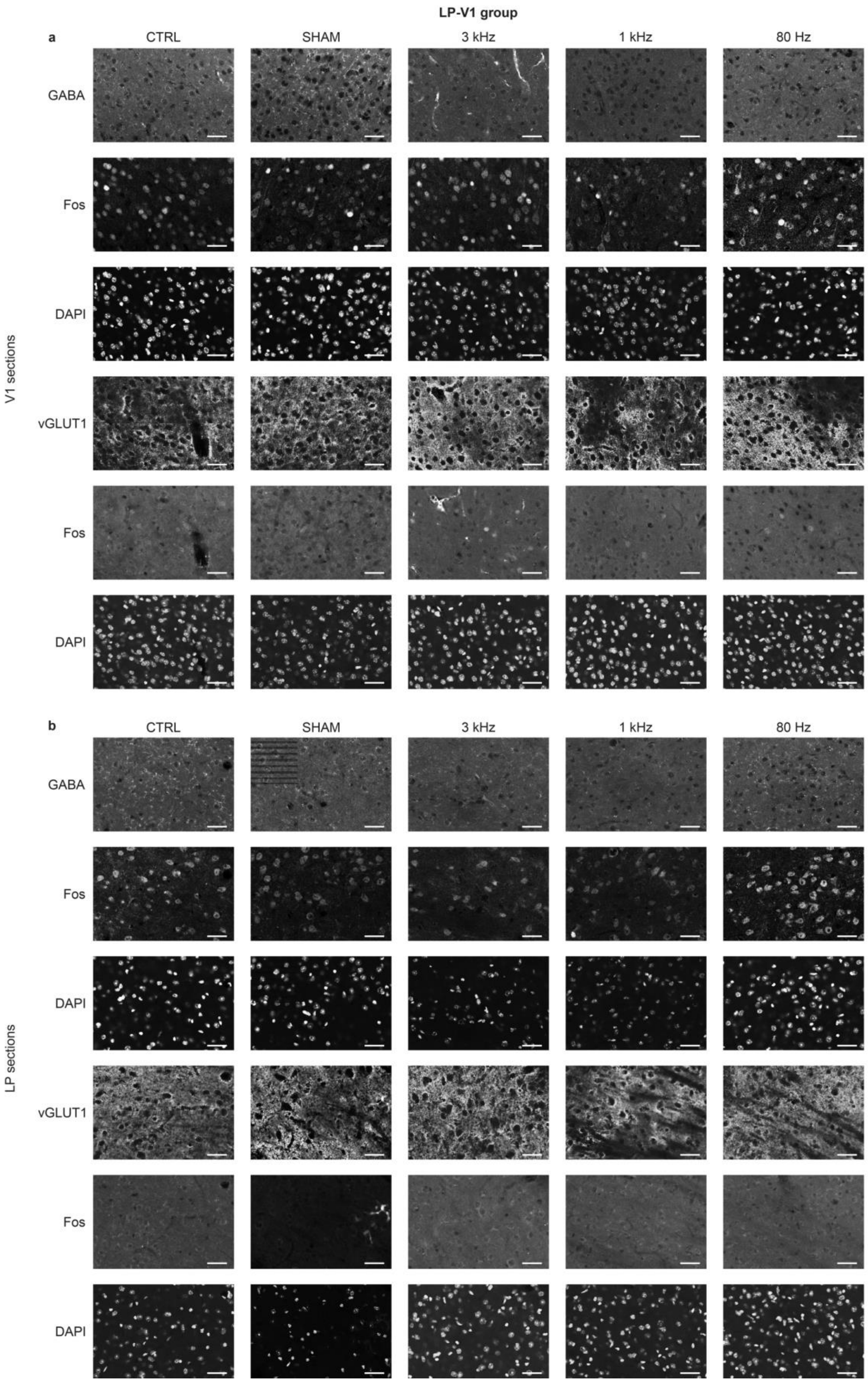

LP-V1 group
a
CTRL
SHAM
3 kHz
1 kHz
80 Hz
V1 sections
GABA
Fos
DAPI
vGLUT1
Fos
DAPI
b
CTRL
SHAM
3 kHz
1 kHz
80 Hz
LP sections
GABA
Fos
DAPI
vGLUT1
Fos
DAPI

*Supplementary Fig. 17 | PRF-dependent activation of glutamatergic and GABAergic neurons for the LP-V1 group.* ***a****, Representative IHC images of GABA and vGLUT1 co-stained with Fos in the V1 for CTRL, SHAM, 3 kHz, 1 kHz, and 80 Hz conditions. Scale bar, 50 µm.* ***b****, Representative IHC images of GABA and vGLUT1 co-stained with Fos in the LP for CTRL, SHAM, 3 kHz, 1 kHz, and 80 Hz conditions. Scale bar, 50 µm.*

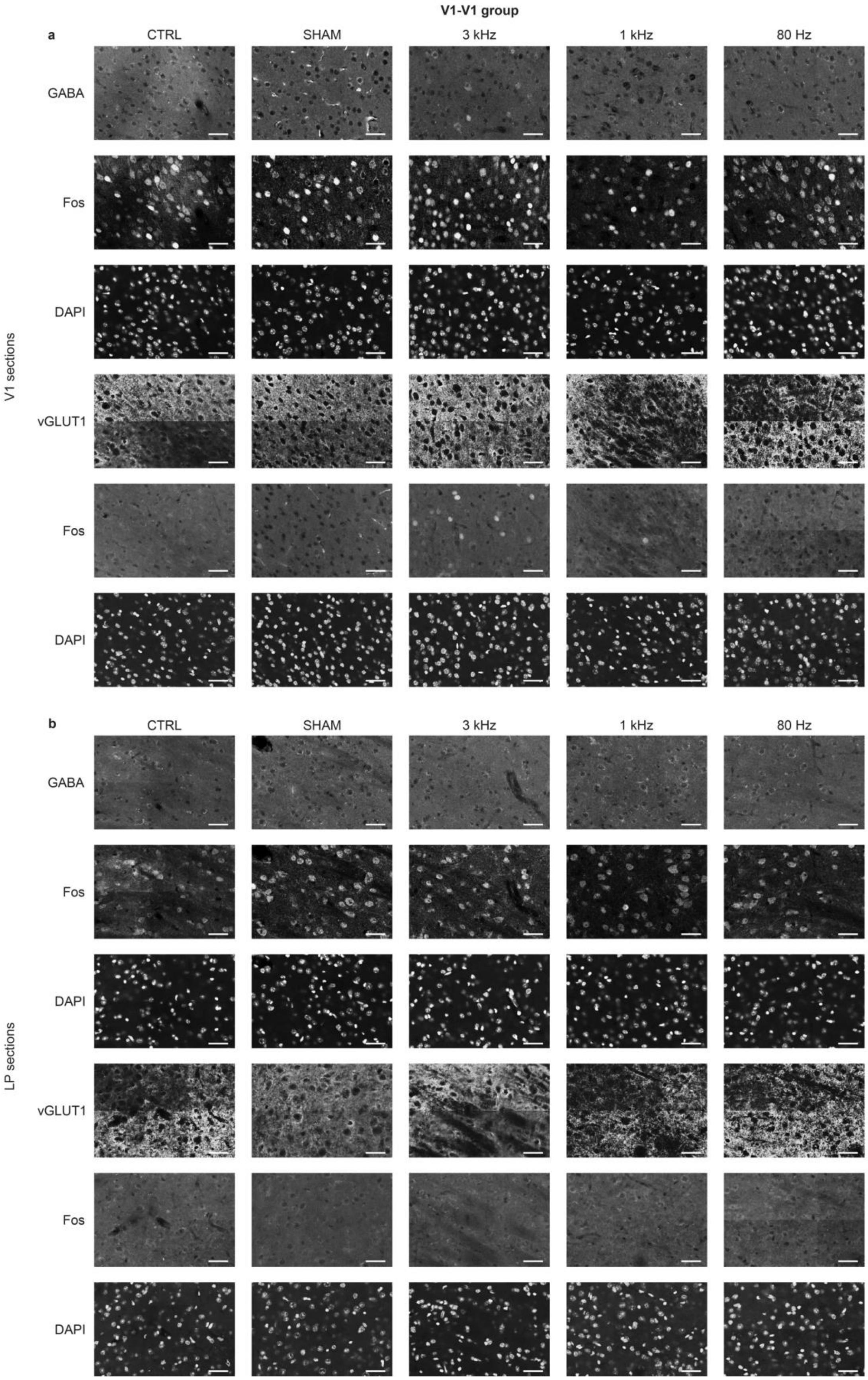
V1-V1 group
a
CTRL
SHAM
3 kHz
1 kHz
80 Hz
GABA
Fos
DAPI
vGLUT1
Fos
DAPI
V1 sections
b
CTRL
SHAM
3 kHz
1 kHz
80 Hz
GABA
Fos
DAPI
vGLUT1
Fos
DAPI
LP sections

*Supplementary Fig. 18 | PRF-dependent activation of glutamatergic and GABAergic neurons for the V1-V1 group.* ***a****, Representative IHC images of GABA and vGLUT1 co-stained with Fos in the V1 for CTRL, SHAM, 3 kHz, 1 kHz, and 80 Hz conditions. Scale bar, 50 µm.* ***b****, Representative IHC images of GABA and vGLUT1 co-stained with Fos in the LP for CTRL, SHAM, 3 kHz, 1 kHz, and 80 Hz conditions. Scale bar, 50 µm.*
